# Crossovers are regulated by a conserved and disordered synaptonemal complex domain

**DOI:** 10.1101/2024.08.05.605830

**Authors:** Ana Rita Rodrigues Neves, Ivana Čavka, Tobias Rausch, Simone Köhler

## Abstract

To ensure the accurate segregation of homologous chromosomes and enhance the genetic diversity in the progeny, meiosis depends on the formation of crossovers between homologous chromosomes. The number and distribution of these crossovers must be precisely regulated through crossover assurance and interference to prevent chromosome missegregation and genomic instability. Here we show that the regulation of crossovers depends on a disordered domain within the synaptonemal complex, which is highly conserved. This domain is located at the C-terminus of the central element protein SYP-4 in *C. elegans*. While not necessary for synapsis, the C-terminus of SYP-4 is crucial for both crossover assurance and interference. Although the SYP-4 C-terminus contains many potential phosphorylation sites, we found that phosphorylation is not the primary regulator of crossover events. Instead, we discovered that nine conserved phenylalanines recruit a pro-crossover factor predicted to be an E3 ligase and regulate the physical properties of the synaptonemal complex. We propose that this conserved and disordered domain plays a crucial role in maintaining the synaptonemal complex in an activated state to promote crossing-over. This activation allows the synaptonemal complex to regulate the number and distribution of crossovers along chromosomes, thereby protecting the genome for future generations.

## Introduction

Meiosis is crucial for sexual reproduction since it segregates not only sister chromatids but also homologous chromosomes to generate gametes with decreased ploidy reducing the chromosome count by half. Errors in the segregation of homologous chromosomes are a major cause of aneuploidy, which can lead to infertility, congenital conditions and miscarriages (1; 2).

The successful separation of homologous chromosomes during meiosis I depends on the formation of crossovers between the homologs. Crossovers (COs) not only increase the genetic diversity of the gametes but also create a physical link between the chromosomes known as chiasma. These chiasmata hold the homologs together until anaphase I and in their absence homologous chromosome segregation is impaired (3; 4; 5). The formation of COs is initiated by the introduction of DNA double-strand breaks (DSBs) that are repaired by homologous recombination. However, only a small subset of DSBs is processed into actual COs as an excessive number of COs can be deleterious (6; 7). Three primary mechanisms regulate crossover formation in meiosis: crossover assurance ensures that each pair of homologs receives at least one obligate CO; crossover interference establishes a non-random distribution of COs, with neighbouring COs spaced further apart than expected by chance; and crossover homeostasis maintains a constant number of COs within each meiotic nucleus to safeguard the system against deficiencies or surpluses of DSBs (8). These highly regulated - or class I - crossovers require ZMM proteins which are named for their members in budding yeast Zip1/2/3/4, Msh4/5, and Mer3. Additionally, many organisms may also acquire a small number of non-interfering class II COs that are formed independently of the class I CO pathway and may function as an alternative repair pathway for surplus DSBs (9; 10; 11). While the regulation of essential class I COs through assurance, interference, and homeostasis is highly conserved across the tree of life, the precise mechanisms by which CO formation is controlled remain unclear.

Specifically, the role of the synaptonemal complex (SC) in regulating crossover formation is a topic of ongoing debate. Through the process of synapsis, this protein structure brings the paired homologous chromosomes into close proximity of around 100 nm (12; 13). The SC is formed by several interacting coiled-coil proteins. In the nematode *Caenorhabditis elegans*, the SC consists of eight interdependent proteins: six coiled-coil proteins, SYP-1-6, and two Skp1-related proteins, SKR-1/2 (14; 15; 16; 17; 18; 19; 20). While the SC is crucial for CO formation in most species, its exact role in the precise patterning of class I COs is still ambiguous. In budding yeast, CO patterning is established independently of SC formation, though SC formation is necessary for the completion of CO formation (21; 22). In contrast, COs are formed in the absence of SC in plants, but CO patterning strictly depends on the SC with a complete loss of CO assurance and interference in absence of the SC (23; 24). Similarly, earlier studies in *C. elegans*, where only class I COs are observed under normal circumstances (10), demonstrated that defects in SC assembly by partial depletion or mutation of its components also give rise to reduced interference (25; 26; 18).

The interdependence of SC integrity and CO patterning in plants and *C. elegans* alongside the biophysical properties of the SC have led to the proposal that an SC-dependent ‘coarsening’ mechanism establishes crossover assurance and interference (27; 28; 29; 23; 24; 30; 31). In this model, a pro-crossover factor, presumably the E3 RING ligase ZHP-3 in *C. elegans*, or HEI10 in plants, initially loads along the SC using its liquid-like properties to diffuse along synapsed chromosomes (27; 28). Indeed, recent single molecule tracking experiments directly confirmed the diffusion of ZHP-3 along SCs (32). At the same time, ZHP-3/HEI10 binds to recombination intermediates, undergoing a coarsening process that selects a few widely spaced recombination intermediates to become designated CO sites (29; 30).

The coarsening model accurately accounts for the loss of both assurance and interference when the coarsening factor HEI10 is not confined to SCs along paired homologous chromosomes in synapsis-deficient plant mutants. In such cases, HEI10 is thought to diffuse and coarsen *in trans* across all chromosomes in the nucleus in a dosage-dependent manner, rather than *in cis* along individual chromosomes, resulting in a random distribution of CO events (23; 24; 30; 33; 31). However, the model fails to explain why interference is detected before SC formation in budding yeast (21; 22). Therefore, it remains unclear whether the observed coarsening process is the actual cause or just a consequence of crossover designation, or whether different species use different regulatory mechanisms to limit crossover designation.

Furthermore, directly testing the effect of the SC on crossover regulation remains challenging, since crossover formation depends on synapsis in *C. elegans* and most other model organisms (34; 14; 35; 36). Moreover, the molecular mechanism by which the potential coarsening factor is recruited to the SC remains unknown.

In this study, we show that the last 114 amino acids of SYP-4 are crucial for regulating crossover formation during meiosis in *C. elegans*. Deleting this region or mutating conserved phenylalanines allows for synapsis but disrupts both crossover assurance and interference, resulting in increased embryonic lethality. We identify a direct link between these crossover defects and the failure to localise ZHP-3 to the SC, accompanied by changes in the biophysical properties of the SC. Our findings indicate that the C-terminus of SYP-4 recruits ZHP-3 to the SC and modulates its biophysical characteristics, which is essential for proper crossover regulation. The function of the C-terminus of SYP-4 as a critical regulator of crossover formation is likely conserved across species.

## Results

Our preliminary data indicated that a frameshift mutation within the last 19 amino acids combined with the addition of a 3X Flag tag at the C-terminus of SYP-4 constitutes a separation-of-function allele that supports synapsis but causes a severe reduction in crossover interference strength during meiosis I (37). This observation suggested that the C-terminus of SYP-4 is involved in the regulation of crossover formation. To understand the role of SYP-4 in this process, we first analysed the amino acid sequence conservation of SYP-4 proteins within the *Caenorhabitis* genus (Fig. 1A). We found conserved regions at the N- and C-terminus of SYP-4. The conserved N-terminus of SYP-4 is predicted to form coiled-coil domains that may interact with other SYP proteins within the synaptonemal complex (16; 19). By contrast, the conserved C-terminus of SYP-4 is predicted to be largely disordered and its conservation suggests that it may have an important and conserved role (Fig. 1A). Notably, our previous single-molecule localisation data suggested that the C-terminus of SYP-4 is not embedded within the SC but protrudes above and below the SC in *C. elegans* (Fig. 1B).

**Figure 1:**
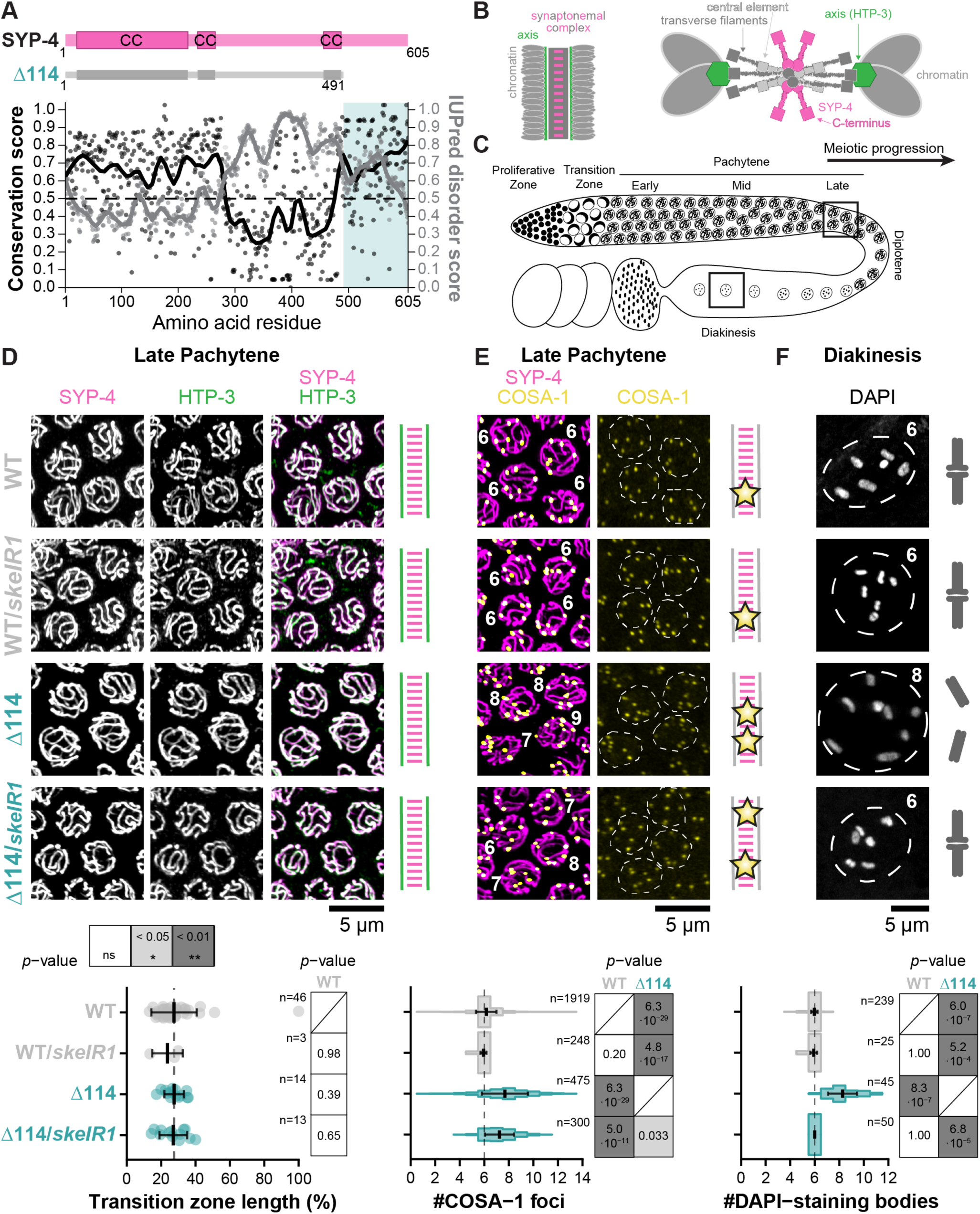
The last 114 amino acids of SYP-4 constitute a highly conserved domain that is dispensable for synapsis but essential for crossover regulation. (A) Both the N- and C-terminus of *C. elegans* SYP-4 are highly conserved (black filled circles; smoothed black line). While the N-terminus is predicted to fold into a coiled-coil (CC) structure, the C-terminus is predicted to be disordered (gray filled circles; smoothed gray line). To test the functionality of this conserved but disordered C-terminal region, we generated the truncated *syp-4*^Δ*114*^ allele using genome editing. Cartoons of SYP-4 are shown on top. (B) Single-molecule localisation data predicts the C-terminus of SYP-4 (magenta squares) in the central element of the SC in frontal view (left) and above and below the synaptonemal complex in cross-sectional view (right). Cartoons are based on data from (37). (C) The diagram of a *C. elegans* germline depicts the progression through the meiotic prophase I stages along the gonad. Images in D and E were acquired in late pachytene, and images in F were acquired in diakinesis (black boxes). (D) Maximum intensity projections of late pachytene nuclei stained for the HA-tagged synaptonemal complex protein SYP-4 (magenta, left) and the axis protein HTP-3 (green, center). The merged image is shown on the right. Cartoons illustrate the major finding of complete synapsis across all genotypes with perfect co-localisation of axes (green) and SYP-4 (magenta). The time required for synapsis quantified as transition zone length (bottom) from initiation of synapsis to completion of synapsis is not changed in homozygous *syp-4*^Δ*114*^ or heterozygous balanced *syp-4*^Δ*114*^/*skeIR1* animals compared to WT animals. The vertical dashed line shows the average transition zone length in WT animals for comparison. Error bars show mean ± standard deviations. *P*-values were calculated using the Mann-Whitney *U* test and corrected using the Benjamini-Hochberg method. (E) Maximum intensity projections of late pachytene nuclei stained for the HA-tagged synaptonemal complex protein SYP-4 (magenta) and the Halo-tagged crossover marker COSA-1 (yellow). Cartoons highlight that crossover interference may be lost or reduced in homozygous and heterozygous *syp-4*^Δ*114*^ animals where some chromosomes acquire more than one COSA-1 focus (stars). Quantification of COSA-1 foci in late pachytene nuclei (bottom) shows an increase in the number of foci in both homozygous *syp-4*^Δ*114*^ and heterozygous balanced *syp-4*^Δ*114*^/*skeIR1* animals compared to WT animals suggesting a semi-dominant effect of the SYP-4 C-terminal truncation. The vertical dashed line at 6 COSA-1 foci shows the expected number of COSA-1 foci in late pachytene nuclei in WT. Error bars show mean ± standard deviations. *P*-values were calculated using a Gamma-Poisson generalised linear model and corrected using the Benjamini-Hochberg method. (F) Maximum intensity projections of diakinesis nuclei, counterstained with DAPI, show some homolog pairs separating into univalents in *syp-4*^Δ*114*^ animals (cartoons). Quantification is shown at the bottom. The vertical dashed line at 6 DAPI-staining bodies corresponds to the expected number of DAPI-staining bodies per diakinesis nuclei in WT. Error bars show mean ± standard deviations. *P*-values were calculated using a Gamma-Poisson generalised linear model and corrected using the Benjamini-Hochberg method.

### The conserved C-terminus of SYP-4 is dispensable for synapsis

To understand the role of the C-terminus of SYP-4 in meiosis, we generated a C-terminally truncated SYP-4 allele, in which the last 114 amino acids were deleted (Fig. 1A, Δ114). In *C. elegans*, the fidelity of chromosome segregation during the meiotic divisions can readily be assessed by quantifying the embryonic lethality and incidence of male progeny: mutations that impair the segregation of chromosomes in meiosis will produce aneuploid progeny giving rise to high embryonic lethality if any of the autosomes is missegregated or male offspring if the X chromosome is missegregated. The transgenic *syp-4*^Δ*114*^ animals showed both high lethality and high incidence of males compared to *syp-4^wt^* animals (Fig. S1, 90.52 ± 2.98% vs −2.38 ± 6.07% and 39.74 ± 10.46% vs 0.07 ± 0.16%, respectively), suggesting that chromosomes were missegregated during meiosis.

The low viability of these animals required us to maintain *syp-4*^Δ*114*^ animals as heterozygous animals in the presence of a wild-type *syp-4* allele. To this end, we first used the *hT2* balancer chromosome consisting of a balanced translocation between chromosomes I and III (38). However, balanced *syp-4*^Δ*114*^/*hT2* animals still showed an increase in lethality and incidence of male progeny compared to *syp-4^wt^*/*hT2* animals (Fig. S2, 85.61 ± 5.14% vs 76.12 ± 1.62% - a 75% embryonic lethality is expected as a result of the lethal *let-?(q782)* mutation and segregation of the balancer chromosomes - and 4.57 ± 5.27% vs 0.22 ± 0.37%, respectively) suggesting that the *syp-4*^Δ*114*^ mutation is (semi-)dominant. Additionally, we frequently observed the acquisition of a *syp-4* wild-type allele on the *syp-4*^Δ*114*^ allele-containing chromosome (Fig. S2C, P1 and P2 lane) indicating that non-homologous recombination occurred in the balanced strain. Therefore, we generated a *syp-4*-specific balancer by introgressing the *ske61* allele (Fig. S2D), which replaces the essential gene *let-383* by a GFP transgene, from the Hawaiian into the Bristol background. This *skeIR1* balancer fully rescued the embryonic lethality and incidence of male progeny of the *syp-4*^Δ*114*^ allele presenting levels comparable to *syp-4^wt^*/*skeIR1* balanced animals (Fig. S1, 25.23 ± 8.08% vs 24.75 ± 5.71% and, 0.37 ± 0.56% vs 0%, respectively).

Mutations in genes constituting the synaptonemal complex (SC) typically cause defects in the assembly of the SC which can drastically reduce the fertility of such animals (34; 14; 15; 16; 25; 26; 18; 19; 39; 20). We therefore tested whether the reduced fertility of *syp-4*^Δ*114*^ animals was caused by defects in SC assembly. To this end, we assessed the co-localisation of an axis protein, HTP-3, and the SC component SYP-4 during pachytene. In both, wild-type and *syp-4*^Δ*114*^ animals, axes and SC were perfectly co-localised along the entire length of chromosomes in pachytene suggesting that the *syp-4*^Δ*114*^ allele supports complete synapsis (Fig. 1C,D). We also measured the length of the transition zone corresponding to the leptotene and zygotene stages of meiosis as a marker for the duration of synapsis (40). The length of the transition zone in *syp-4*^Δ*114*^ animals was indistinguishable from wild-type animals (Fig. 1D, 27.52 ± 5.66% vs 27.49 ± 13.19%; Fig. S3A). Measuring the expression levels and loading of SYP-4 on the synaptonemal complex using Western Blotting and quantitative imaging, respectively (Fig. S3B-D) revealed that SYP-4^Δ114^ is more abundant than wild-type SYP-4, and the timing of SYP-4 loading on the chromosome axes is not altered in absence of the C-terminus of SYP-4 in *syp-4*^Δ*114*^ animals (Fig. S3B-D). These results show that the C-terminus of SYP-4 is dispensable for the assembly and maintenance of the SC but is required for other meiotic processes.

### The C-terminus of SYP-4 regulates crossover patterning

The synaptonemal complex is not only essential for maintaining the pairing of homologous chromosomes but also for crossover formation (25; 26; 23; 24; 37). We therefore next investigated crossover formation in *syp-4*^Δ*114*^ animals. In *C. elegans*, the crossover factor COSA-1 acts as a cytological marker for designated crossover sites in late pachytene (41). We visualised COSA-1 using an endogenously engineered HaloTag, and quantified the number of COSA-1 foci per nucleus in late pachytene (Fig. 1E). We automated the quantification of COSA-1 foci by integrating automated nucleus segmentation (42) with spot identification using spotMAX (Padovani *et al*, manuscript in preparation). A comparison between our automated analysis and manual quantifications for 481 nuclei representative of the dataset demonstrated that the automated pipeline yielded results consistent with manual counts (Fig. S4). Using the automated quantification of COSA-1 foci, we found that the number of foci per nucleus was slightly elevated from 6.16 ± 0.84 in wild-type to 7.68 ± 1.86 in *syp-4*^Δ*114*^ animals (Fig. 1E). Interestingly, the number of COSA-1 foci was also elevated in heterozygous *syp-4*^Δ*114*^/*skeIR1* animals (7.24 ± 1.18) suggesting a (semi-)dominant effect of the *syp-4*^Δ*114*^ allele. However, while the number of COSA-1 foci in heterozygous *syp-4*^Δ*114*^/*skeIR1* animals resembled the number in homozygous *syp-4*^Δ*114*^ animals, the elevated embryonic lethality and the increase in male progeny was fully rescued in heterozygous animals compared to homozygous *syp-4*^Δ*114*^ animals (Fig.S1). Thus, the slight increase in COSA-1 foci cannot explain the drastic fertility defects observed in *syp-4*^Δ*114*^ animals consistent with previous findings showing that additional crossovers rarely give rise to missegregation of chromosomes in meiosis (7).

While COSA-1 reliably marks designated crossover sites in late pachytene in *C. elegans* (41), such designated sites must then progress to form chiasmata to establish the bivalent structure in diakinesis. In wild-type animals, the presence of six bright COSA-1 foci correlates with the formation of six chiasmata, which organise the six pairs of chromosomes into six bivalents observed as 5.97 ± 0.26 DAPI-staining bodies in diakiness (Fig. 1C,F). Similarly, heterozygous *syp-4^wt^*/*skeIR1* and *syp-4*^Δ*114*^/*skeIR1* animals exhibited six DAPI-staining bodies (5.92 ± 0.28 and 6.00 ± 0.00, respectively), indicating the establishment of at least one chiasma per homologous chromosome pair. However, in *syp-4*^Δ*114*^ animals, we observed 8.27 ± 1.18 DAPI-staining bodies suggesting that on average only 4 chromosomes typically form bivalents, while two pairs of homologs remain as four univalents (Fig. 1F). This finding implies that not all designated crossovers mature into chiasmata in *syp-4*^Δ*114*^ animals. Alternatively, it suggests that some chromosomes may acquire multiple crossovers while others lack crossovers entirely. Indeed, we observed that both *syp-4*^Δ*114*^ and *syp-4*^Δ*114*^/*skeIR1* animals display ring-shaped bivalents, indicating the presence of two chiasmata on these chromosomes suggesting that some chromosomes may receive multiple crossovers (Fig. S5A,B). To further characterise the defects in crossover regulation in *syp-4*^Δ*114*^ animals, we traced single chromosomes within late pachytene nuclei and quantified the number of COSA-1 foci on each chromosome (Fig. S6A,B). Our analysis revealed that 32% corresponding to approximately 2 out of 6 chromosomes were devoid of COSA-1 foci, mirroring the proportion of homologous chromosome pairs that fail to form bivalents (Fig. 1F). Additionally, more than 33% of chromosomes in *syp-4*^Δ*114*^ late pachytene nuclei displayed multiple COSA-1 foci. To assess the strength of crossover interference within these chromosomes, we fitted a gamma distribution to the normalised inter-COSA-1 focus distances. This analysis allowed us to estimate the shape parameter (γ), serving as a relative measure of crossover interference strength. In this context, a γ value of 1 denotes no interference, while higher shape parameters indicates stronger interference (43; 44; 11). In our analysis, we included four wild-type nuclei, each containing a rare occurrence of one chromosome bearing two COSA-1 foci as a proxy for wild-type interference. While these WT chromosomes exhibited robust crossover interference (γ=42.9) consistent with prior studies (26), *syp-4*^Δ*114*^ animals displayed a significantly reduced interference with a γ=2.7 (Fig. S6C). Together, these data indicate a substantial decline in both crossover assurance and interference in *syp-4*^Δ*114*^ animals.

Thus, the C-terminus of SYP-4 is essential to regulate crossover formation to ensure that each pair of chromosomes receives exactly one crossover.

### Putative phosphorylation sites within the C-terminus of SYP-4 do not play a major role in crossover patterning

Phosphorylations play a critical role in regulating many aspects during meiotic progression (45). In *C. elegans*, the phosphorylation of SYP-4 regulates DNA double-strand break (DSB) formation and the choice of the recombination pathway (46; 47). Consequently, we hypothesised that the C-terminus of SYP-4 might undergo phosphorylations to control crossover formation. To identify potential phosphorylation sites in the C-terminus of SYP-4 systematically, we conducted mass-spectrometry analysis of SYP-4 purified from young adults (Table S1). We identified four potential sites: S447, S485, S496, and S554. Notably, phosphorylation of S447 was previously linked to directing DSB repair toward homologous recombination, thereby preventing repair via the non-homologous end joining pathway (47). However, mutating S447 alone did not affect crossover interference and assurance, with each chromosome still acquiring exactly one crossover in both phosphodead and phosphomimetic S447 mutants (47). We therefore explored whether multiple phosphorylations of the C-terminus of SYP-4 might collaborate to robustly regulate crossover numbers. To test this hypothesis, we mutated all four identified sites from our mass-spectrometry analysis, along with six additional Ser/Thr residues that were not covered or under-represented in our mass-spectrometry analysis (Table S1), resulting in the *syp-4^10SD^*/*syp-4^10SA^* phosphomimetic/phosphodead mutants (Fig. 2A, top). Both *syp-4^10SD^* and *syp-4^10SA^* mutants showed slightly reduced fertility with a small increase in embryonic lethality and higher numbers of male progeny (Fig. S1, 8.49 ± 4.71% and 5.86 ± 16.03%, and 2.83 ± 1.32% and 1.13 ± 0.76%, respectively) indicating that some aspects of meiosis may be impaired. However, both phosphomimetic and phosphodead *syp-4^10SD^* and *syp-4^10SA^* animals exhibited normal synapsis (Fig. 2B, Fig. S3A, and S7A) and displayed only mild defects in crossover regulation, characterised by a slight increase in COSA-1 foci from 6.16 ± 0.84 in wild-type animals to 6.78 ± 1.15 and 6.34 ± 1.15 in *syp-4^10SD^* and *syp-4^10SA^*animals, respectively (Fig. 2C, and S7B). Crossover assurance was also not affected in *syp-4^10SD^* or *syp-4^10SA^* animals and all chromosomes received at least one crossover giving rise to six DAPI-staining bodies (Fig. 2D, and S7C).

**Figure 2:**
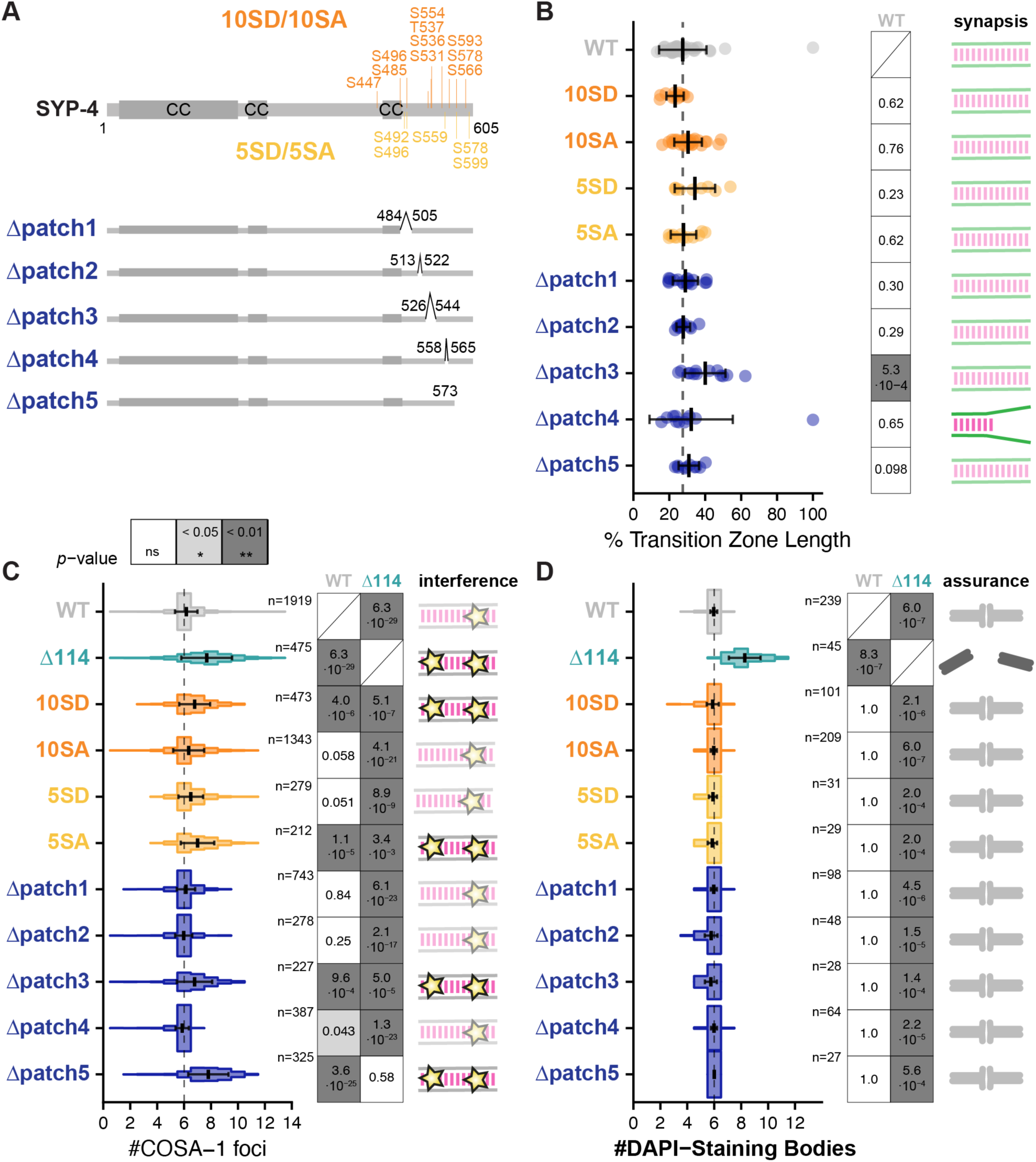
Phosphorylation of the C-terminus of SYP-4 and the last 32 amino acids of SYP-4 modulate crossover interference but are dispensable for crossover assurance. (A) Diagram of SYP-4 indicating the serine/threonine residues substituted by aspartate residues in SYP-4 10SD and 5SD phosphomimetic, or by alanine residues in SYP-4 10SA and 5SA phosphodead mutants (top), and the short internal/terminal deletions within the C-terminus of SYP-4 (bottom). CC corresponds to predicted coiled-coil domains. (B) The length of transition zone is unaffected in all SYP-4 phosphomutants and almost all short deletions compared to WT animals but it is slightly extended in Δpatch3 animals as indicated by the cartoons (right). The vertical dashed line corresponds to the average transition zone length observed in WT animals. Error bars show mean ± standard deviations. *P*-values were calculated using the Mann-Whitney *U* test and corrected using the Benjamini-Hochberg method. (C) The quantification of COSA-1 foci in late pachytene nuclei shows a slight increase in the number of foci for all phosphomutants, Δpatch3 and Δpatch5 animals compared to WT animals. Moreover, the observed increase of COSA-1 foci in Δpatch5 animals is similar to the increase observed in *syp-4*^Δ^*^114^* animals. The vertical dashed line at 6 COSA-1 foci corresponds to the expected number of COSA-1 foci per nucleus in WT. Cartoons depict the likely decrease in interference observed for some of the mutants. Error bars show mean ± standard deviations. *P*-values were calculated using a Gamma-Poisson generalised linear model and corrected using the Benjamini-Hochberg method. (D) Quantification of DAPI-staining bodies in diakinesis nuclei indicates the presence of 6 bivalents in most of the quantified nuclei in phosphomutant, short deletion, and WT animals. Cartoons depict the robust bivalent formation in all mutants. Error bars show mean ± standard deviations. *P*-values were calculated using a Gamma-Poisson generalised linear model and corrected using the Benjamini-Hochberg method.

In parallel, we performed an *in silico* analysis using the Eukaryotic Linear Motif (ELM) resource to predict motifs in the C-terminus of SYP-4 (48). This analysis revealed five potential binding sites for BRC-1, which localises to the synaptonemal complex and is implicated in regulating the choice of pathways during meiotic DSB repair (49; 50; 51; 52; 53). To investigate, whether binding of BRC-1 to the C-terminus of SYP-4 contributes to crossover regulation, we generated phosphomimetic (*syp-4^5SD^*) and phosphodead (*syp-4^5SA^*) mutants (Fig. 2A). Both mutants exhibited almost normal fertility (Fig. S1) and normal synaptonemal complex assembly and maintenance (Fig. 2B, Fig. S3A, and S7A). Consistent with their largely normal fertility, both *syp-4^5SD^*and *syp-4^5SA^* animals also showed robust crossover regulation, with only a slight increase in designated crossover sites marked by Halo-tagged COSA-1 in the phosphodead *syp-4^5SA^* mutant compared to the wild type (Fig. 2C, and S7B; 7.00 ± 1.24 vs 6.16 ± 0.84). The number of DAPI-staining bodies remained unaffected by both mutations (Fig. 2D, and S7C).

Together, these findings indicate that phosphorylations of the C-terminus of SYP-4 are dispensable for synapsis and crossover assurance but influence the recombination landscape in *C. elegans*, as previously proposed (46; 47). Phosphorylation of potential BRC-1 motifs in the C-terminus of SYP-4 also modulate crossover patterning, but they are not essential for ensuring robust crossover assurance and interference. Similarly, prior research indicated that BRC-1 monitors and modulates meiotic recombination (49; 50; 51; 52; 53). However, the mutations targeting potential phosphorylation sites did not replicate the significant defects in crossover regulation seen in *syp-4*^Δ*114*^ animals.

### The last 32 amino acids of SYP-4 ensure robust crossover interference

To investigate whether specific regions are required for regulating crossover formation in *C. elegans*, we identified five conserved patches within the C-terminus of SYP-4 (Fig. 2A, bottom). Deletion of any of these patches resulted in viable offspring (Fig. S1). The embryonic lethality and male progeny of *syp-4*^Δ*patch1*^, *syp-4*^Δ*patch2*^, *syp-4*^Δ*patch3*^, and *syp-4*^Δ*patch4*^ animals closely resembled wild-type animals, but *syp-4*^Δ*patch5*^ animals exhibited significantly higher embryonic lethality and a higher percentage of male progeny (Fig. S1, 38 ± 6.88% and 5.28 ± 2.13%, respectively), albeit lower than in *syp-4*^Δ*114*^ animals. All animals successfully formed the synaptonemal complex (Fig. S7A), and the length of the transition zone resembled its length in wild-types animals for four of the five strains carrying the small deletions. However, *syp-4*^Δ*patch3*^ animals exhibited a slight extension of the transition zone length compared to wild-type animals (39.97 ± 11.28% vs 27.49 ± 13.19%; Fig. 2B and S3A). Given that the deletion of the last 114 amino acids of SYP-4 in *syp-4*^Δ*114*^ animals does not cause synapsis defects, we speculate that the extension of transition zone in *syp-4*^Δ*patch3*^ animals may result from defective folding of SYP-4 and/or reduced protein stability, rather than patch3 region being essential for SC assembly.

We further assessed whether deleting short stretches in the C-terminus of SYP-4 affects crossover regulation by quantifying COSA-1 foci in late pachytene. While *syp-4*^Δ*patch1*^, *syp-4*^Δ*patch2*^, and *syp-4*^Δ*patch4*^ animals displayed an average of six COSA-1 foci, similar to wild-type, *syp-4*^Δ*patch3*^ and *syp-4*^Δ*patch5*^ animals exhibited an increase to 6.78 ± 1.31 and 7.80 ± 1.49 foci, respectively (Fig. 2C, and S7B). The slight rise in the number of COSA-1 foci observed in *syp-4*^Δ*patch3*^ animals likely stems from delayed synaptonemal complex assembly, consistent with previous results (25; 26; 54). However, the more drastic increase in *syp-4*^Δ*patch5*^ animals mirrors the defects seen in *syp-4*^Δ*114*^ animals where 7.68 ± 1.89 COSA-1 foci were observed (Fig. 2C). Despite these variations in the number of COSA-1 foci, all animals with deletions in the C-terminus of SYP-4 exhibited six DAPI-staining bodies in diakinesis, indicating complete crossover assurance (Fig. 2D, and S7C).

Together, these findings suggest that the last 32 amino acids constituting patch5 are essential for restricting the number of designated crossover sites to exactly six sites per nucleus. These results are also consistent with our preliminary data showing that a frameshift mutation affecting the last 19 amino acids in conjunction with the insertion of a 3X Flag epitope tag (*syp-4*^CmutFlag^) severely impairs crossover interference but not assurance (37).

### Conserved phenylalanines within the C-terminus of SYP-4 are involved in crossover patterning

Since neither phosphorylations nor single conserved patches in the C-terminus of SYP-4 could explain the severe defects in crossover regulation observed in *syp-4*^Δ*114*^ animals, we investigated other characteristics of the C-terminus of SYP-4 to determine its role in crossover regulation. Analysing the amino acid composition revealed that the C-terminus of SYP-4 is enriched in phenylalanines: while Phe constitutes only 4% of all residues in the full-length protein, Phe makes up 14% of the C-terminus of SYP-4. Nine of the sixteen Phe in the C-terminus of SYP-4 are located in conserved motifs resembling nematode-specific LIR-motifs that typically mediate the interaction with Atg8-related proteins in autophagy (48). To test whether these Phe have a specific function during meiosis in SYP-4, we generated a mutant, *syp-4^9FA^*, replacing the nine Phe in LIR-motifs by alanines (Fig. 3A). *syp-4^9FA^* exhibited very high embryonic lethality and produced a high number of male progeny (Fig. S1, 91.75 ± 6.36% and 20.12 ± 13.65%, respectively) that resembled the brood counts of *syp-4*^Δ*114*^ animals. Like *syp-4*^Δ*114*^ animals, *syp-4^9FA^* animals were also synapsis proficient (Fig. 3B,C), although the expression level of *syp-4^9FA^* was lower than for wild-type SYP-4 and *syp-4*^Δ*114*^ (Fig. S3B-D). In accordance with the strong reduction in fertility, *syp-4^9FA^* animals also displayed a slight increase in the number of COSA-1 foci to 7.59 ± 1.88 compared to 6.16 ± 0.84 in wild-type animals, which resembles the increase in *syp-4*^Δ*114*^ animals (Fig. 3D,E). We also observed a slight increase in the number of COSA-1 foci in balanced heterozygous *syp-4^9FA^*/*skeIR1* animals to 7.08 ± 1.19 foci suggesting that the *syp-4^9FA^* mutation is also semi-dominant (Fig. 3D,E). Similar to *syp-4*^Δ*114*^ animals, *syp-4^9FA^* animals but not balanced *syp-4^9FA^*/*skeIR1* showed an increase in the number of DAPI-staining bodies suggesting that not all homologs are held together by crossover events (7.17 ± 0.83, Fig. 3F,G). Indeed, COSA-1 foci are not evenly distributed across chromosomes and some chromosomes in *syp-4^9FA^* animals lack any COSA-1 focus while other chromosomes have more than one focus (Fig. S6A,B). We also observe ring-shaped bivalents in diakinesis in *syp-4^9FA^* animals suggesting the presence of at least two crossovers on these chromosomes (Fig. S5).

**Figure 3:**
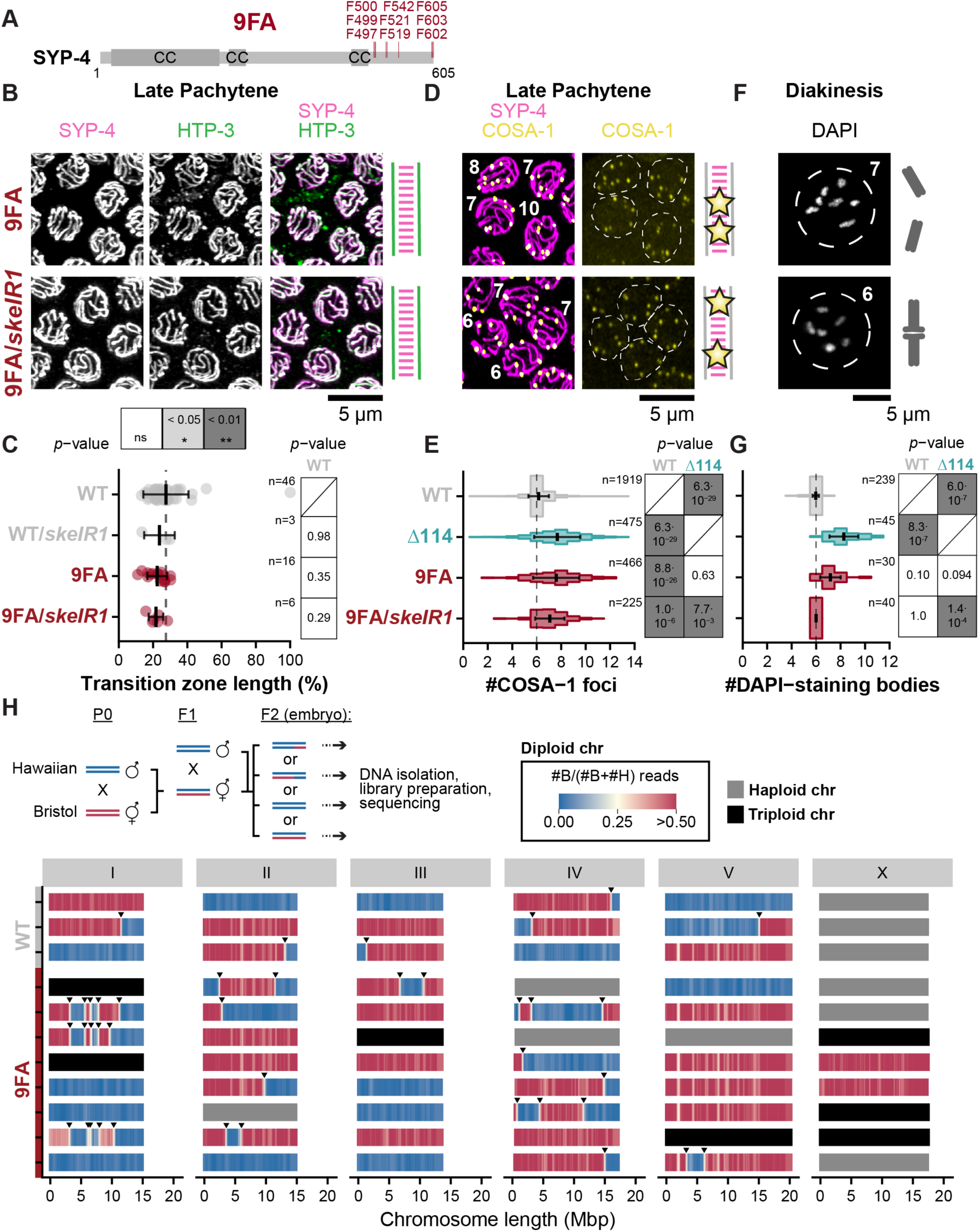
The C-terminus of SYP-4 regulates crossover formation via conserved phenylalanine residues. (A) Diagram of SYP-4 indicating the phenylalanine residues substituted by alanine residues in *syp-4^9FA^* animals. CC corresponds to predicted coiled-coil domains. (B) Maximum intensity projections of late pachytene nuclei stained for the HA-tagged synaptonemal complex protein SYP-4 (magenta, left) and the axis protein HTP-3 (green, center). The merged image is shown on the right. (C) The transition zone length is not altered in homozygous *syp-4^9FA^* or heterozygous balanced *syp-4^9FA^*/*skeIR1* animals compared to WT animals. The vertical dashed line corresponds to the average transition zone length observed in WT animals. Error bars show mean ± standard deviations. *P*-values were calculated using the Mann-Whitney *U* test and corrected using the Benjamini-Hochberg method. (D) Maximum intensity projections of late pachytene nuclei stained for the HA-tagged synaptonemal complex protein SYP-4 (magenta) and the Halo-tagged crossover marker COSA-1 (yellow). (E) Quantification of COSA-1 foci in late pachytene nuclei shows an increase in the number of foci in both homozygous *syp-4^9FA^*and heterozygous balanced *syp-4^9FA^*/*skeIR1* animals compared to WT animals suggesting a semi-dominant effect of the *syp-4^9FA^* allele. The vertical dashed line at 6 COSA-1 foci shows the expected number of COSA-1 foci in late pachytene nuclei in WT. *P*-values were calculated using a Gamma-Poisson generalised linear model and corrected using the Benjamini-Hochberg method. (G) Maximum intensity projections of diakinesis nuclei counterstained with DAPI. (H) Quantification of DAPI-staining bodies in diakinesis nuclei shows an increase in the number of DAPI-staining bodies in homozygous *syp-4^9FA^* animals compared to WT indicating the presence of univalents, while the number of DAPI-staining bodies is not increased in heterozygous balanced *syp-4^9FA^*/*skeIR1* animals. *P*-values were calculated using a Gamma-Poisson generalised linear model and corrected using the Benjamini-Hochberg method. (H) Genetic mapping of crossover sites confirms loss of assurance and interference in *syp-4^9FA^* animals. A schematic representation of the experimental strategy followed to genetically map crossover sites is shown on top. Chromosome tracks show the ratio between Bristol-specific reads and the total number of specific reads along the chromosomes. A ratio of zero (blue) corresponds to homozygous regions derived from the Hawaiian background whereas a ratio of 0.5 or more (red) corresponds to heterozygous regions containing both Hawaiian- and Bristol-specific sequences. The transition from homozygous (blue) to heterozygous (red) regions corresponds to crossover sites (arrow heads). Chromosomes that were found to contain a single copy (haploid, gray) or three copies (triploid, black) based on the average copy number per chromosome were not considered for the detection of crossover sites.

In summary, both *syp-4*^Δ*114*^ and *syp-4^9FA^* animals exhibit defects in the distribution of crossover events with defects in both crossover assurance and interference. Indeed, the distribution of COSA-1 foci along synaptonemal complexes resembles a random gamma distribution in both *syp-4*^Δ*114*^ and *syp-4^9FA^* animals suggesting that interference is strongly impaired (Fig. S6C). However, the intensity of COSA-1 foci is more variable in *syp-4*^Δ*114*^ and *syp-4^9FA^* animals than in wild-type animals (Fig. S6D). Since only “bright” COSA-1 foci typically mature into crossovers (55), we tested whether all COSA-1 foci give rise to genetic crossovers in *syp-4^9FA^* animals by mapping recombination events in hybrids of two divergent *C. elegans* strains, Bristol N2 and Hawaiian CB4856, carrying a *syp-4^wt^* or a *syp-4^9FA^* allele using whole genome sequencing (Fig. 3H). We analysed recombination events in genomes of backcrossed F2 embryos rather than adult F2 and their progeny to account for the high embryonic lethality of *syp-4^9FA^*animals (Fig. 3H). As expected, about 50% of the X chromosomes were haploid indicating male embryos. However, we also found triploid X chromosomes in *syp-4^9FA^* animals indicating that the X chromosome may also be missegregated. We therefore excluded the X chromosome from all further analysis. In wild-type animals, all autosomes were diploid and we identified six crossovers in 15 autosomes suggesting that 40% of the chromosomes received a crossover (Fig. 3H). This value is slightly lower than the expected 50% of chromosomes stemming from the fact that only one crossover is generated for each chromosome with two sister chromatids. While the slightly lower frequency of COs per chromosome compared to expected values may be due to the low number of chromosomes analysed, we also cannot exclude that the relatively low resolution achievable by single-embryo whole genome sequencing prevents us from detecting all crossovers. In *syp-4^9FA^* animals, 20% of autosomes were aneuploid suggesting that autosomes were also missegregated as already indicated by the increased number of DAPI-staining bodies (Fig. 3H). Similarly, our analysis revealed the presence of multiple crossovers at variable distances in *syp-4^9FA^* animals. However, we found several identical recombination sites on chromosome I in three out of eight *syp-4^9FA^* animals indicating recombination scars from the introgression of the *skeIR1* balancer from the Hawaiian background (see methods). We therefore also excluded chromosome I from our analysis. On the remaining autosomes, we found on average 0.73 crossovers corresponding to 1.46 COSA-1 foci per pair of homologous chromosomes (n=26), suggesting that interference is reduced. Indeed, the distribution of genetically mapped crossovers closely followed the distribution of COSA-1 foci with a shape factor of 2.7 confirming the strong decrease in interference in *syp-4^9FA^* animals compared to wild-type animals (Fig. S6C). Thus, the aberrant localisation of COSA-1 in *syp-4^9FA^* animals closely resembles the genetic outcome we observe by recombination mapping.

### The C-terminus of SYP-4 recruits ZHP-3 to the synaptonemal complex

Although the SC is pivotal for crossover formation in most species, including *C. elegans*, crossovers still occur in synapsis-deficient *Arabidopsis* meiocytes (23; 24; 56). However, these crossovers are no longer regulated through crossover assurance and interference but are distributed randomly along chromosomes. This lack of regulation was attributed to the failure to recruit the E3 ligase pro-crossover factor HEI10 due to defects in SC assembly (23; 24; 31).

Given the resemblance of the strong defects in crossover regulation observed in *syp-4*^Δ*114*^ and *syp-4^9FA^* animals to the phenotype in SC-deficient plants, we investigated whether the HEI10-homolog ZHP-3 is similarly mislocalised in the C-terminally mutated SYP-4 animals. While ZHP-3 co-localises with SC proteins to polycomplexes in early transition zone or along the SC until mid-pachytene in WT animals, it is predominantly nucleoplasmic in *syp-4*^Δ*114*^ and *syp-4^9FA^* animals (Fig. 4A, and Fig. S8A).

**Figure 4:**
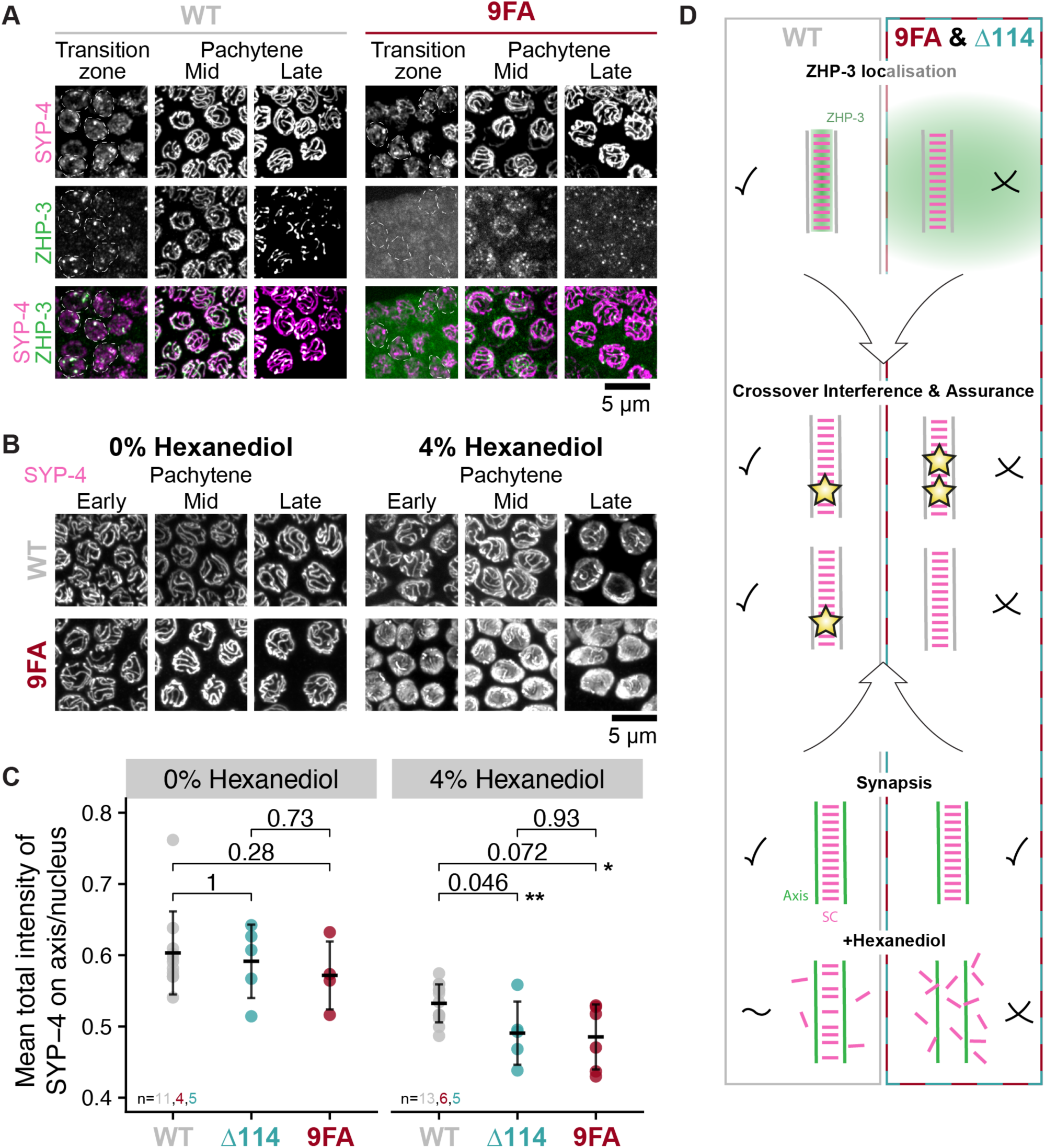
The defects in crossover regulation in *syp-4^9FA^* and *syp-4*^Δ*114*^ animals are accompanied by a mislocalisation of ZHP-3 and changes in the biophysical properties of the SC. (A) Maximum intensity projections of transition zone, mid and late pachytene nuclei stained for HA-tagged SYP-4 (magenta, top) and V5-tagged ZHP-3 (green, middle) show that the co-localisation of ZHP-3 with polycomplexes (transition zone) or SCs (pachytene) is lost in *syp-4^9FA^* animals. The merged image is shown on the right. Nuclei containing poly-complexes during transition zone are encircled with a dashed line. (B) Maximum intensity projections of early, mid and late pachytene nuclei from extruded gonads treated with 0% *w/v*) or 4% (*w/v*) 1,6-hexanediol and stained for HA-tagged synaptonemal complex protein SYP-4 (magenta, top) and the axis protein HTP-3 (green, middle). The merged image is shown at the bottom. (C) SCs in *syp-4^9FA^* and *syp-4*^Δ*114*^ animals are more sensitive to 1,6-hexanediol than SCs of WT animals. The ratio of the total SYP-4 intensity on the axis to total SYP-4 intensity per nucleus in WT (gray), *syp-4^9FA^* (red) and *syp-4*^Δ*114*^ (cyan) are shown. In untreated pachytene nuclei, the majority of SYP-4 signal is found on the axis (axis/nucleus ratio > 0.5), while SYP-4 is more nucleoplasmic in presence of 4% (*w/v*) 1,6-hexanediol. The reduction of SYP-4 on the axis is higher in *syp-4^9FA^* and *syp-4*^Δ*114*^ animals. Average axis/nucleus ratios during pachytene. Error bars show mean ± standard deviations. *P*-values were calculated using the Mann-Whitney *U* test (* *p*-value<0.1, ** *p*<0.05). (D) Diagram summarising the role of nine conserved phenylalanine residues within the C-terminus of SYP-4 in ZHP3 localisation (top) and SC stability (bottom) to regulate crossing-over (center).

In contrast, most ZHP-3 foci co-localise with COSA-1 at the end of pachytene, likely indicating crossover sites in both WT and *syp-4*^Δ*114*^ or *syp-4^9FA^* animals (Fig. 4A, and Fig. S8A). This finding suggests that phenylalanines within the C-terminus of SYP-4 recruit the pro-crossover factor ZHP-3 to the SC. Interestingly, a non-synonymous single nucleotide variation switching a leucine to a phenylalanine in the C-terminus of the human central element protein SIX6OS1 (57) was associated with an increased recombination rate in women (58). This finding implies that the function of the disordered C-terminus of the central element protein SYP-4 in *C. elegans* may be conserved across animals.

The synaptonemal complex has previously been demonstrated to exhibit liquid-like properties (27). Given that the accumulation of hydrophobic phenylalanines in disordered protein domains has been described to enhance the tendency of such domains to undergo phase separation (59; 60; 61; 62; 63), we investigated whether the critical phenylalanines we identified affect the biophysical characteristics of the SC. To do so, we examined the response of SCs from both WT and *syp-4*^Δ*114*^ or *syp-4^9FA^* animals to hexanediol. In the absence of hexanediol, SC proteins assemble along chromosome axes in both WT and *syp-4*^Δ*114*^ or *syp-4^9FA^* SCs. However, in the presence of 4% hexanediol, the SC partially dissolves into the nucleoplasm (Fig. 4B,C and Fig. S8B,C) (27). Notably, while approximately 53% of SYP-4 remains associated with the axis in WT SCs under these conditions, only 49% of SYP-4^Δ114^ or SYP-4^9FA^ remains bound. This observation suggests that the phenylalanines located in the C-terminus of SYP-4 play a role in stabilising the SC, thereby influencing its biophysical properties.

Together, our data demonstrates a conserved role of the disordered C-terminus of the central element protein SYP-4 in meiotic crossover regulation, revealing that phenylalanines within this region are essential for recruiting the pro-crossover factor ZHP-3 and stabilising the SC.

## Discussion

Our study revealed that the C-terminus of SYP-4 is dispensable for synaptonemal complex assembly, but it plays a crucial role in controlling the formation of crossovers, regulating both crossover interference and crossover assurance.

Specifically, we demonstrated that removing the C-terminus of SYP-4, or mutating nine phenylalanines within this region, not only alters the physical characteristics of the SC but also prevents the binding of the pro-crossover factor and E3 ligase ZHP-3 in both *syp-4*^Δ*114*^ and *syp-4^9FA^* animals. This misplacement of ZHP-3 from the SC to the nucleoplasm resulted in significant impairments in crossover assurance and interference, resembling the crossover defects observed in plants deficient in synapsis (23; 24; 56).

In plants, this finding prompted the hypothesis that a primary role of the SC is to restrict the movement of the pro-crossover factor HEI10, the plant homolog of ZHP-3. With restricted movement, recombination events along a single SC must compete for a limited pool of HEI10 molecules, leading to the formation of only a few crossovers that are spaced far apart — a process termed “coarsening” (30). Therefore, the absence of synapsis in plants, or in our case, the failure to recruit ZHP-3 to the SC due to phenylalanine mutations in the C-terminus of SYP-4, may replace the localised coarsening within the SC with a broader coarsening across the nucleus (31). This mislocalisation of ZHP-3/HEI10 may result in a loss of crossover interference and assurance, while maintaining a relatively stable number of crossovers through crossover homeostasis (31).

Alternatively, ZHP-3 might have additional crucial roles at the synaptonemal complex in regulating crossover formation. Evidence suggests that the localisation of ZHP-3 and its heterodimer partner, ZHP-4, is intricately linked to their enzymatic activity, as mutations in their E3 RING finger domains not only abolish crossover formation but also prevent their loading along the SC (64). Therefore, it is plausible that the enzymatic activity of the ZHP-3/ZHP-4 heterodimer may serve additional functions at the SC that are independent of and may precede their coarsening behaviour.

While the entire disordered C-terminus of SYP-4, or the phenylalanines dispersed within it, are crucial for establishing both crossover assurance and interference, we also identified a specific conserved segment within this domain that is essential for ensuring robust crossover interference. This segment contains the final 32 amino acids of the C-terminus and is essential for ensuring that crossover interference operates across distances longer than individual chromosomes in *C. elegans*. This finding confirms our earlier observations, where preliminary data indicated that a mutation in the last 19 amino acids of SYP-4 significantly diminished the strength of crossover interference (37).

Surprisingly, despite the abundance of potential phosphorylation sites within the C-terminus of SYP-4, phosphorylations are not a major regulatory mechanism for CO formation but they make CO regulation more resilient. Our findings are thus in agreement with prior data showing that recombination is influenced by SYP-4 phosphorylation, with post-translational modifications within its C-terminus playing a role in determining the recombination pathway and maintaining SC integrity under conditions of elevated DNA double-strand breaks (46; 47). Thus, post-translational modifications might serve to increase the robustness of crossover regulation in *C. elegans*, particularly under challenging circumstances.

The high conservation of the SYP-4 C-terminus suggests that its function in regulating crossover events may be conserved in other nematodes and maybe even in more distant metazoans. For example, a mutation in the likely disordered C-terminal domain in the mammalian *SIX6OS1* central element component was also linked to defects in crossover regulation (58; 57). These data therefore suggest that the role of the SC, and in particular, of the disordered C-terminus of SYP-4, in regulating crossover formation may be conserved across metazoans.

## Materials and Methods

### Maintenance and culture conditions of *C. elegans* strains

All *C. elegans* strains were cultured at 20°C under standard conditions on nematode growth medium plates inoculated with *E. coli* OP50 (65). All experiments were performed using young adults 18-24 hours post-L4. For experiments with balanced strains, we selected non-green fluorescent homozygous L4-staged animals. A list of all strains used in this study can be found in Table S2.

### Generation of *C. elegans* strains by CRISPR/Cas9-mediated genome editing

New alleles in this study were generated by CRISPR/Cas9-mediated genome editing by injection of preassembled Cas9-gRNA RNP complexes as described previously (66). The Cas9-NLS protein was purchased either from the EMBL Protein Expression and Purification Core Facility or from IDT. The common tracrRNA as well as all crRNAs listed in Table S3 were purchased from IDT. Wild-type sequences were obtained from WormBase (67) and modified to yield the repair template sequences used to generate each new allele as listed in Table S4. For the *ske19* and *ske71* alleles, short ssDNA oligos (Sigma-Aldrich) were used as repair templates. For the *ske25* allele, a biotinylated-dsDNA obtained by PCR from a gBlock (IDT) was used as a repair template. For *ske61* allele, a dsDNA was obtained by PCR from pCFJ1415 (68). For all remaining alleles the repair templates were obtained by PCR from gBlocks (IDT). To enrich for successfully injected progeny, two plasmids carrying red fluorescent transgenes (2.5 ng/µL pCFJ90 [Pmyo-2::mCherry] and 5 ng/µL pCFJ104 [Pmyo-3::mCherry] (69)) were co-injected with the Cas9-gRNA RNP and repair template. Individual F1s expressing either of the two plasmids were picked onto new plates and were screened for correctly edited genes by PCR two or more days later (Table S5). For alleles containing silent mutations in *syp-4*, we introduced or removed restriction sites to distinguish mutant from wild-type alleles (Table S5). All sequences were confirmed by Sanger sequencing (Eurofins Genomics). All epitope-tagged *syp-4*, *zhp-3* and *cosa-1* wild-type genes generated in this study were functional with no major increases in embryonic lethality or percentage of male progeny (Table S6). Strains containing new alleles were outcrossed at least twice, or experiments were performed using at least two synonymous transgenic lines derived from different CRISPR/Cas9-editing events. To generate a balancer for *syp-4*, the essential gene *let-383* was replaced by a codon optimized GFP transgene in the Hawaiian CB4856 background. This allele was then introgressed into the Bristol N2 background by nine consecutive crosses.

### Viability Assays

To determine the viability of eggs, L4 hermaphrodites were singled onto individual plates to lay eggs. After each laying period of 12-24 hours, the number of fertilised eggs was counted, and the individual worms were transferred to new plates. This process was repeated until they stopped laying eggs. The surviving progeny was counted once they reached the adult stage.

### SYP-4 multiple sequence alignment, amino acid conservation and disorder prediction

We searched for homologs of the *C. elegans* SYP-4 protein sequence (uniprot ID Q9N5K3) in all proteomes listed in Table S7 using a protein hidden markov model (HMM) (phmmer). The hits obtained from this first search were used to perform a reciprocal mapping by searching them against the *C. elegans* proteome. Only hits that successfully mapped back to *C. elegans* SYP-4 were then used to perform a multiple sequence alignment (MSA) using MAFFT (Multiple Alignment using Fast Fourier Transform) with the genafpair algorithm and a maximum of 1000 iterations (70). Both amino acid conservation scores and protein disorder prediction were retrieved from JalView (71). The conservation scores correspond to the calculated quality scores which are based on BLOSUM62 substitution matrix. Protein disorder scores are based on IUPred prediction for short regions. PairCoil2 (72) was used to predict coiled-coil motifs in SYP-4 protein sequence (P-score>0.025).

### Immunofluorescence

Animals were dissected 24 hours post-L4 and stained as described previously in (73) with modifications described in (74) and mounted in ProLong Glass antifade mounting medium (Invitrogen, P36984).

The following primary antibodies were used: chicken anti-HTP-3 (1:500, (75)), mouse anti-HA (1:1250, Invitrogen, #A190-138A), or mouse anti-V5 (1:500, Invitrogen, #R960-25). As secondary antibodies the following were used: Alexa Fluor 488 donkey anti-chicken (1:500, Jackson ImmunoResearch, 703-545-155), Alexa Fluor 546 goat anti-chicken (1:500, Invitrogen, #A-11040), Alexa Fluor 546 donkey anti-mouse (1:500, Invitrogen, #A10036), Alexa Fluor 647 donkey anti-mouse (1:500, Jackson ImmunoResearch, 715-605-150). Sample blocking and antibody incubation were performed in 1X Roche blocking solution in PBST.

To visualise COSA-1 foci, animals carrying a HaloTag::*cosa-1* allele from well-fed plates were incubated overnight with 6.67 µM JF669 HaloTag ligand (Lavis Lab, JBG-27-075A) or 1.3 µM JF646 HaloTag ligand (Promega, GA1120) in a 20 µL droplet of M9 buffer containing *E. coli* OP-50 and 0.2% (v/v) Tween-20. To assess the loading of SYP-4 throughout meiosis and test the sensitivity of SYP-4 to 4% (v/v) 1,6-hexanediol, both *syp-4* mutant animals carrying the endogenously tagged HaloTag::*cosa-1* allele and control animals with untagged *cosa-1* were fed overnight on plates seeded with OP50 *E. coli* in a 20 µL drop of 6.67 µM JF669 HaloTag ligand (Lavis Lab, JBG-27-075A) in M9 buffer with 0.2% (v/v) Tween-20 and dissected together on the same slide the next day. For hexanediol treatments, the dissected samples were incubated for 30 s with 4% (v/v) 1,6-hexanediol before fixation as described in (27).

Images were acquired within two weeks after mounting. At least two independent immunofluorescence experiments were performed.

### Fluorescence Microscopy

For quantification of COSA-1 foci and DAPI-staining bodies, 0.16 µm-spaced *z*-stacks were acquired on a Zeiss LSM 880 Airyscan laser scanning confocal microscope with a 63X 1.4 NA DIC M27 oil plan-apochromat objective using the FastAiryScan mode. The images were deconvolved using the 3D auto mode from the Airyscan processing tool available in the Zen Black v14.0.15.201 software (Zeiss).

For all other experiments, 0.16-0.20 µm-spaced *z*-stacks were acquired on an Olympus spinning disk confocal system IXplore SpinSR with a 60X 1.4 NA oil plan-apochromat objective using the cellSens software (Olympus). For visualization of ZHP-3, quantification of SYP-4 loading throughout pachytene and hexanediol experiments images were acquired using the higher resolution SoRa disk with a 3.2X magnification. For transition zone quantification, images were acquired using a 50 µm disk with a 1X magnification.

All image manipulations, such as maximum intensity projection, tile stitching, channel intensity adjustment, and colour correction were performed using the free software Fiji (76).

### Image quantification

#### Segmentation of meiotic nuclei

DAPI-stained meiotic nuclei were blurred with a 2.5 blurring factor and segmented in 3D using a custom pipeline based on the Cellpose2.0 neural network (77) using a custom model (https://github.com/KoehlerLab/Cellpose_germlineNuclei/blob/main/Cellpose_germlineNucleicellpose_germlineNuclei_KoehlerLab) as described in (42). To remove incomplete nuclei, segmented nuclei masks touching the first or last slice in *z* were disregarded.

#### Quantification of SYP-4 loading throughout meiosis and SYP-4 sensitivity to 1,6-hexanediol

We used custom Python and R scripts to quantify the loading of SYP-4 along the axis or the sensitivity of SYP-4 to 1,6-hexanediol (https://github.com/KoehlerLab/SYPquant/tree/main/SYPquant/SCintensityAnalysis). To calculate the background signal of SYP-4::HA labelling, we calculated the average pixel intensity within 10 and 50 pixels from the segmented objects (including filtered out objects). To quantify the SYP-4::HA signal along the chromosome axes, we segmented the HTP-3 signal within each segmented nucleus using the mean thresholding method from the Python library scikit-image v0.19.1 (78). For each nucleus, the total intensities of SYP-4::HA on the HTP-3 mask and in the nucleus were calculated after subtracting the average background signal. For the quantification of SYP-4::HA loading on the axis, the total intensity of SYP-4::HA on the axis was normalised to the average total intensity of SYP-4::HA signal on the axis in wild-type samples from the same slide. For the quantification of SYP-4::HA sensitivity to 1,6-hexanediol, we calculated the ratio between the intensity of SYP-4::HA on the axis and in the nucleus.

A region of interest was manually defined for each germline and objects outside this region were removed. To remove potential fragmented meiotic nuclei, apoptotic nuclei or somatic nuclei, only segmented nuclei with a volume between 10 and 60 µm^3^ (excluding), a sphericity higher than 0.4 and an intensity ratio between 0.3 and 0.8 (including) were considered. To assess the loading and sensitivity of SYP-4 along the pachytene region, the germlines were re-oriented perpendicularly to the manually annotated end of transition zone from left to right. The germlines were straightened by applying a Loess function to the centroid positions of all segmented nuclei. The germline length was normalized from the beginning of the transition zone or beginning of pachytene to the end of pachytene to assess SYP-4::HA loading and effects of hexanediol treatment, respectively, using manual annotations. This region was then divided into 11 bins and averages of the normalised intensity or intensity ratio and standard deviations were calculated for each bin. To assess the average sensitivity of SYP-4 to hexanediol treatment, the average of the ratio of SYP-4 total intensity on axis and total nucleus over the bins was calculated per gonad.

#### Quantification of COSA-1

For the quantification of COSA-1 foci, well segmented nuclei in late pachytene were selected manually. We then used spotMAX v0.8.0, a 3D gaussian fitter software (Padovani *et al.*, manuscript in preparation), to quantify the number of foci per segmented meiotic nucleus. The foci were initially thresholded inside the segmentation masks using the Otsu thresholding method and fit by a 3D gaussian function.

##### Chromosome tracing

The segmented nuclei were first cropped using the object’s bounding box. Each of the six chromosome filaments were semi-manually traced using the Fiji plugin BigTrace (v.0.05 or v.0.2.1) on semi-automate mode (https://github.com/ekatrukha/BigTrace) based on SYP-4::HA signal. For visualisation purposes, traced chromosomes were straightened using an 11-pixel wide line around the center line using BigTrace (v.0.2.1) and the respective maximum intensity projection of the merged SYP-4 and COSA-1 channels was generated.

##### COSA-1 localisation along the chromosomes

The COSA-1 centroid position was mapped to the position along the traced chromosome using a custom Python script (https://github.com/KoehlerLab/SYPquant/tree/main/SYPquant/ ChrTraceAnalysis) that comprises the following steps: (i) calculation of the offset between the COSA-1 and SYP-4 channel to correct the COSA-1 centroid coordinates; (ii) spline fitting of BigTrace’s output (trace point coordinates) and sampling of equidistant points along the interpolated spline; (iii) mapping of the COSA-1 centroid positions onto the closest trace corresponding to the point on the trace with the smallest eucledian distance to the COSA-1 centroid position.

#### Quantification of DAPI-staining bodies

The number of DAPI-staining bodies in diakinesis was quantified manually in 3D stacks from DAPI-counterstained samples using Fiji.

#### Quantification of transition zone length

Transition zone was quantified manually in maximum intensity projections using Fiji. Transition zone was defined from the first nuclei with visible HTP-3 stretches until the last row of nuclei with incomplete synapsis based on HTP-3 and SYP-4::HA immunofluorescence signal localisation.

### Recombination mapping

#### Sample Preparation

To map crossover frequencies genetically, L4-staged hermaphrodites of different genotypes in the wild-type Bristol background (wild-type N2 or *syp-4(9FA)::ha (ske39-1)/skeIR1* I; *halo::cosa-1 (ske25)* III) were crossed to males of the same genotype in the Hawaiian background (wild-type CB4856 and *syp-4(9FA)::ha (ske39-2)/ske61* I, respectively) as illustrated in Fig. 3H (P0: Bristol background in red, Hawaiian background in blue). The F1 hermaphrodite Bristol/Hawaiian progeny (N2/CB4856 or *syp-4(9FA)::ha (ske39-1)*/*syp-4(9FA)::ha (ske39-2)* I; *halo::cosa-1 (ske25)*/+ III) was crossed with male homozygous CB4856 Hawaiian animals, and 24 hours later hermaphrodite animals were transferred to empty 6 cm nematode growth medium agar plates containing a 20 µL drop of 10 mM serotonin [Sigma-Aldrich, H7752] within a palmitic acid ring to increase the rate of egg laying and prevent worms from leaving the agar plate, respectively (79; 80). Single F2 embryos were collected for a period of 6 hours followed by DNA isolation, library preparation and whole genome sequencing. Singled F2 embryos were transferred onto lysis buffer (1x DreamTaq Buffer [ThermoScientific, B65], 1 mg/mL Proteinase K [EMBL Protein Expression and Purification Core Facility]) and stored at −80°C for at least 24 hours. The genomic DNA was isolated after a 1 hour incubation at 65°C followed by Proteinase K inactivation at 95°C for 15 min. The genomic DNA was amplified using the PicoPLEX® Single Cell WGA Kit (TAKARA, R300672) according to the manufacturer’s instructions. The amplified genomic DNA was purified using a 1.8x volume of SPRIselect beads (Beckman Coulter, B23319) followed by Tn5-based tagmentation and 300PE Illumina sequencing on an Illumina NextSeq 2000 platform (EMBL Gene Core Facility) to a final coverage of 1.8-6.7X per sample.

### Data processing

The sequencing quality was assessed using FastQC (https://www.bioinformatics.babraham.ac.uk/projects/fastqc/) and MultiQC (81). The adapter sequences from Illumina index primers were removed using Trimmomatic (82). Reads mapping to the standard-8 database (10/9/2023) available at https://benlangmead.github.io/aws-indexes/k2 were filtered out using Kraken2 (83). The remaining reads were mapped to the softmasked N2 Bristol (BioProject PRJNA13758) and CB4856 Hawaiian (BioProject PRJNA275000) WS288 genome releases available on WormBase (https://wormbase.org) (67) using BWA-MEM (84). Read sorting, duplicate removal and read filtering were performed using SAMtools (85). Reads with a mapping quality score below ten and mitochondrial reads were not considered.

### Data analysis

To identify crossover sites genetically, we counted the number of reads in 5 kbp windows that mapped exclusively to the N2 Bristol or CB4856 Hawaiian genome, respectively, using SAMtools (85) and Alfred (86). To infer genotypes, we calculated the gliding average of the ratio between Bristol-specific reads and the total number of Bristol- and Hawaiian-specific reads across 100 5 kbp windows. If this ratio was greater than 0.9, we classified it as homozygous Bristol (only observed for haploid chromosomes), and if it was less than 0.25, we classified it as homozygous Hawaiian. Ratios in between indicated a heterozygous genotype. We identified crossovers as points where there was a change in genotype. We only considered transitions supported by more than 1500 reads, with consistent genotypes on either side for at least 15 kbp. Each crossover site was also manually verified.

Recombination sites on chromosome I were excluded from further analysis due to the specific recombination events observed in *syp-4(9FA)::ha (ske39-1)*/*syp-4(9FA)::ha (ske39-2)* I; *halo::cosa-1 (ske25)*/+ III (9FA samples) indicating a mix of Hawaiian and Bristol backgrounds in the parental strain that stems from the introgression of the *skeIR1* allele.

To determine chromosome ploidy, we conducted a copy number variant analysis on 50 kbp windows of reads mapped to the CB4856 Hawaiian WS288 genome using the delly software (86). We created a mappability map following established protocols as described in https://github.com/dellytools/delly using dicey, samtools and BWA (87; 85; 84). Chromosomes with an average copy number falling between 1.45 and 2.5 were classified as diploid, those below 1.45 as haploid, and those above 2.5 as triploid. Only diploid chromosomes were considered for the identification of crossover sites.

### Crossover interference analysis

Crossover interference was measured based on the gamma shape factor of the gamma distribution fitted to the inter-crossover distances for genomic data or inter-COSA-1 distances for cytological data (44). A best-fit gamma shape factor of 1 corresponds to no interference and higher/smaller shape factor values correspond to increasingly positive/negative interference.

### Immunoblotting

To compare SYP-4 protein levels in *syp-4* mutant animals, 20 young adults (24-hour post-L4) were picked into a 30 µL droplet of M9 buffer (22 mM KH_2_PO_4_, 49 mM Na_2_HPO_4_, 86 mM NaCl, 10 mM NH_4_Cl) with 0.1% (v/v) Tween-20 and washed 3 times with M9 buffer with 0.1% (v/v) Tween-20 in 0.2 mL tubes to remove bacteria. Samples were frozen and stored at −80°C for at least 24 hours. Samples were thawed and resuspended in 10 µL benzonase buffer (50 mM Tris-HCl pH 8.0, 2 mM MgCl_2_) with EDTA-free protease inhibitor (Roche, 11873580001, 1 tablet per 25 mL benzonase buffer). To release chromatin-bound proteins and reduce sample viscosity, samples were sonicated in a water bath at 4°C for 5 cycles (sonication cycle: 30 sec ON/ 30 sec OFF) followed by benzonase treament at 37°C for at least 1 hour with 5 U of benzonase (Millipore, 71205). Sample loading buffer with reducing agent (Invitrogen, NP0007, 4X NuPAGE™ LDS Sample Buffer, and NP0004, 10X NuPAGE Sample Reducing Agent) was added to a final 1.5x concentration in a volume of 20 µL. Samples were boiled for 10 min at 70°C to and loaded on a 1.0 mm NuPage precast 4-12% (v/v) Bis-Tris gel (Invitrogen). SDS-PAGE and semi-wet transfer to a methanol-pre-activated polyvinylidene difluoride (PVDF) membrane were performed in an Electrophoresis Mini Gel Tank (Life Technologies, A25977) according to the manufacturer’s instructions. After protein transfer, the PVDF membrane was blocked for 30 min at room temperature in 1X Roche Blocking buffer (Roche, 11096176001) in PBST. The following primary antibodies were used: mouse anti-HA (1:5000, Invitrogen, #A190-138A) and mouse anti-tubulin (1:5000, Sigma-Aldrich, T6199-25UL). Goat anti-mouse-horseradish peroxidase conjugate (1:5000, BioRad, #1706516) was used as secondary antibody. Proteins were detected by chemiluminescence using the Pierce ECL Plus Western Blotting Substrate kit (ThermoScientific, 32132) according to the manufacturer’s instructions and images were acquired on a BioRad system. Image processing and quantification were performed using the free software Fiji (76).

### Phosphoproteomic analysis of SYP-4

#### Meiotic nuclei-enriched sample preparation

A population of *syp-4::ha(ie29); gfp::cosa-1(meIs8)* animals was synchronised by bleaching as described in (88). Around 15 000 synchronised L1-staged worms were plated per 100-mm diameter peptone-enriched nematode growth medium agar plate inoculated with *E. coli* OP-50 and left to grow to young adulthood for 48 hours at 20°C. A total of 10-20 plates were prepared. Meiotic nuclei were isolated in Nuclei Purification Buffer (NPB: 25 mM HEPES pH 7.4, 118 mM NaCl, 48 mM KCl, 2 mM EDTA, 0.5 mM EGTA, 0.2 mM DTT, 0.25 mM spermine, 0.5 mM spermidine, 0.1% (v/v) Tween-20) supplemented with 1X PhosSTOP (Roche, #4906845001) and 2 µg/mL Proteinase Inhibitor (Sigma, #P9599-1ML) based on (89) with some modifications. Briefly, the meiotic nuclei were released from the gonad tissue by dissection with a razor blade in 5 mL NPB buffer in a 60 mm ødish instead of Dounce homogenisation. The broken worm solution was transferred to a pre-chilled 15 mL canonical tube and vortexed on high speed for 30 sec, followed by 5 min on ice. Then, the solution passed through two 40 µm cell strainers (FisherScientific, #07-201-430) and one 20 µm cell strainer (pluriSelect, #43-50020). The isolated nuclei were collected by centrifugation at 2500 x *g* for 5 min at 4°C. The supernatant was removed, the nuclei were resuspended in 200 µL NPB and transferred to a 1.5 mL tube. The meiotic nuclei were spun down at 2500 x *g* for 5 min at 4°C, the supernatant was removed, the nuclei pellet was flash frozen in liquid nitrogen and stored at −80°C until sonication. Three biological replicates were prepared.

#### SYP-4 immunoprecipitation

Isolated meiotic nuclei were resuspended in 500 µL of iCLIP lysis buffer (90) supplemented with 1X PhosSTOP (Roche, #4906845001) and 2 µg/mL Proteinase Inhibitor (Sigma, #P9599-1ML) and incubated on ice for 15 min. Samples were sonicated on a BIORUPTOR® PICO machine (Diagenode) at 4°C in the “low” setting for 5 min (30 sec on; 30 sec off). Samples were treated with 4 U of TURBO DNase (FisherScientific, #10646175) at 37°C for 5 min at 1200 rpm followed by centrifugation at 15 000 x *g* for 15 min at 4°C. The supernatant was transferred to a new tube and 700 µg of protein were incubated with mouse anti-HA antibody-coupled Dynabeads in 500 µL of iCLIP lysis buffer (trypsin-digested samples: 4 µg antibody per 50 µL beads; FisherScientific, #26183 and #10515883; chemotrypsin-digested samples: 10 µg antibody per 1.5 mg beads; FisherScientific, #26183 and ThermoFisher, #14311D). The beads were magnetically separated, washed twice with 900 µL High Salt Wash Buffer, once in 500 µL Wash Buffer (90) and then resuspended in 1X NuPAGE Sample Loading Buffer (Invitrogen, NP0007) and 1X NuPAGE Reducing Agent (Invitrogen, NP0004). Samples were boiled at 95°C for 5 min, loaded on a NuPAGE 4-12% Bis-Tris precast gel (Invitrogen) and ran in 1X NuPAGE MOPS SDS Running Buffer (Invitrogen, NP0001). The gel was stained with Coomassie Staining solution (50% (v/v) EtOH, 10% (v/v) CH_3_COOH, 0.25% (w/v) Coomassie Blue G-250) for 4 hours at room temperature, followed by gel destaining with bidistilled water overnight. The gel was further processed by EMBL Proteomics Core Facility.

#### Mass spectrometry sample preparation

Two biological replicates were digested with trypsin and one with chymotrypsin since trypsin digestion resulted in low coverage of the C-terminus of SYP-4.

Briefly, the bands were cut from the gel and subjected to in-gel digestion with trypsin or chymotrypsin (91). Peptides were extracted from the gel pieces by sonication for 15 min, followed by centrifugation and supernatant collection. A solution of 50:50 water:acetonitrile, 1% (v/v) formic acid (2X the volume of the gel pieces) was added for a second extraction, and the samples were again sonicated for 15 min, centrifuged and the supernatant pooled with the first extract.

#### LC-MS/MS analysis

The pooled supernatants were subjected to speed vacuum centrifugation. The samples were dissolved in 10 µL of reconstitution buffer (96:4 water: acetonitrile, 1% (v/v) formic acid. Samples were injected into an UltiMate 3000 nanoRSLC (Dionex, Sunnydale, CA) coupled with a trapping cartridge (µ-Precolumn C18 PepMap 100, 5µm, 300 µm i.d. x 5 mm, 100 Å) and an analytical column (nanoEase™ M/Z HSS T3 column 75 µm x 250 mm C18, 1.8 µm, 100 Å, Waters). Trapping was carried out with a constant flow of trapping solvent (0.05% (v/v) trifluoroacetic acid in water) at 30 µL/min onto the trapping column for 6 min. Subsequently, peptides were eluted and separated on the analytical column using a gradient composed of Solvent A [(3% (v/v) DMSO, 0.1% (v/v) formic acid in water] and solvent B [3% (v/v) DMSO, 0.1% (v/v) formic acid in acetonitrile] with a constant flow of 0.3 µL/min. During the analytical separation, the percentage of solvent B was stepwise increased from 2% to 4% in 6 min, from 4% to 8% in 1 minute, then 8% to 25% in 41 min, and finally from 25% to 40% in another 5 min. The outlet of the analytical column was coupled directly to an Orbitrap Fusion Lumos (Thermo Scientific, SanJose) mass spectrometer using the nanoFlex source. The peptides were introduced into the Orbitrap Fusion Lumos via a Pico-Tip Emitter 360 µm OD x 20 µm ID; 10 µm tip (New Objective) and an applied spray voltage of 2.4 kV, the instrument was operated in positive mode. The capillary temperature was set at 275°C. Full mass scans were acquired for a mass range of 375-1200 m/z in profile mode in the orbitrap with a resolution of 120000. The filling time was set to a maximum of 50 ms with a limitation of 4e5 ions. The instrument was operated in data-dependent acquisition (DDA) mode, and MSMS scans were acquired in the Orbitrap with a resolution of 30000, a fill time of up to 86 ms and a limitation of 2e5 ions (AGC target). A normalised collision energy of 34 was applied. MS2 data was acquired in profile mode.

#### Identification of phosphorylated sites in SYP-4

Acquired data was processed by IsobarQuant (92) and Mascot (v2.2.07) was used for protein identification. Data was searched against the UniProt *C. elegans* proteome database containing common contaminants, reversed sequences and the sequences of the proteins of interest. The data was searched with the following modifications: Carbamidomethyl (C; fixed modification), Acetyl (N-term), Oxidation (M) and Phospho (STY) (variable modifications). The mass error tolerance for the full scan MS spectra was set to 10 ppm and for the MS/MS spectra to 0.02 Da. A maximum of two missed cleavages was allowed. For protein identification, a minimum of 2 unique peptides with a peptide length of at least seven amino acids and a false discovery rate below 0.01 were required on the peptide and protein level.

### Statistics

All statistical analysis were performed in R. Differences between different strains in embryonic lethality, incidence of male progeny, transition zone length, coefficient of variation of COSA-1 foci intensity and SYP-4 sensitivity to 1,6-hexanediol were tested using the non-parametric Mann-Whitney *U* two-sided test from the R Stats package (v4.3.2). Differences in SYP-4 protein levels quantified by Western Blot were statistically assessed using the non-parametric Wilcoxon Signed-Sum two-sided paired test available in the R Stats package (v4.3.2). To assess whether the automated identification of COSA-1 using spotMAX was statistically different than the manual identification of COSA-1 in the same nucleus a paired two-sided Student’s *t*-test was used available from the R Stats package (v4.3.2). Statistical analysis of the number of COSA-1 foci, number of DAPI-staining bodies and number of ring-shaped DAPI-staining bodies in different strains was performed using a Gamma-Poisson generalised linear model from the bioconductor glmGamPoi package (93), which is less sensitive to sample size differences compared to other tests. When more than 10 comparisons were performed against the same strain a *p*-value correction was performed using the Benjamini-Hochberg correction method available in the R Stats package (v4.3.2).

## Data Availability

Sequencing data for recombination mapping are available on the European Nucleotide Archive under accession number PRJEB76871. All other study data are included in the article and as supplementary information.

## Acknowledgements

We thank the Advanced Light Microscopy Facility (ALMF) at the European Molecular Biology Laboratory (EMBL) in Heidelberg, Evident/Olympus and Zeiss for their support; the Genomics Core Facility at the European Molecular Biology Laboratory (EMBL) in Heidelberg for high-throughput sequencing and expertise; the Proteomics Core Facility at the European Molecular Biology Laboratory (EMBL) for the phosphoproteomics LC-MS/MS sample preparation and analysis; the Protein Expression and Purification Core Facility at the European Molecular Biology Laboratory (EMBL) in Heidelberg for providing purified Cas9-NLS and Proteinase K for this study; IT and HPC resources at the European Molecular Biology Laboratory (EMBL) in Heidelberg for providing essential computational infrastructure; the Media and Lab Kitchen at the European Molecular Biology Laboratory (EMBL) in Heidelberg for providing media, solutions and sterilised material for this study. Some strains were provided by the CGC, which is funded by NIH Office of Research Infrastructure Programs (P40 OD010440). We thank the Dernburg lab (UC Berkeley) for providing the chicken anti-HTP-3 primary antibody; Luke Lavis (Janelia Research Campus) for JF669-HaloTag ligand; Oane Jan Gros for a custom script for cropping individually segmented nuclei for further chromosome tracing; Kausthubh Ramachandran and Gautam Dey for their contribution on SYP-4 MSA; Wolfgang Huber for advise on statistical approaches; Francesco Padovani and Kurt Schmoller for their help with spotMAX. This work was supported by: the European Molecular Biology Laboratory, and by the Deutsche Forschungsgemeinschaft (DFG, German Research Foundation) - project number 452616889.

## References

[1] Hassold T, Hunt P (2001) To err (meiotically) is human: the genesis of human aneuploidy. Nature Reviews Genetics 2(4):280–291.

[2] Nagaoka SI, Hassold TJ, Hunt PA (2012) Human aneuploidy: mechanisms and new insights into an age-old problem. Nature Reviews Genetics 13(7):493–504.

[3] Maguire MP (1974) The need for a chiasma binder. Journal of Theoretical Biology 48(2):485–487.

[4] Zickler D, Kleckner N (1999) Meiotic Chromosomes: Integrating Structure and Function. Annual Review of Genetics 33(1):603–754.

[5] Zickler D, Kleckner N (2015) Recombination, Pairing, and Synapsis of Homologs during Meiosis. Cold Spring Harbor Perspectives in Biology 7(6):a016626.

[6] Serrentino ME, Borde V (2012) The spatial regulation of meiotic recombination hotspots: Are all DSB hotspots crossover hotspots? Experimental Cell Research 318(12):1347–1352.

[7] Hollis JA, et al. (2020) Excess crossovers impede faithful meiotic chromosome segregation in C. elegans. PLOS Genetics 16(9):e1009001.

[8] Wang S, Zickler D, Kleckner N, Zhang L (2015) Meiotic crossover patterns: Obligatory crossover, interference and homeostasis in a single process. Cell Cycle 14(3):305–314.

[9] Lynn A, Soucek R, Börner GV (2007) ZMM proteins during meiosis: Crossover artists at work. Chromosome Research 15(5):591–605.

[10] Saito TT, Colaiácovo MP (2017) Regulation of Crossover Frequency and Distribution during Meiotic Recombination. Cold Spring Harbor Symposia on Quantitative Biology 82(3):223–234.

[11] Girard C, Zwicker D, Mercier R (2023) The regulation of meiotic crossover distribution: a coarse solution to a century-old mystery? Biochemical Society Transactions 51(3):1179–1190.

[12] Moses MJ (1968) Synaptinemal Complex. Annual Review of Genetics 2(1):363–412.

[13] Westergaard M, von Wettstein D (1972) THE SYNAPTINEMAL COMPLEX. Annual Review of Genetics 6(1):71–110.

[14] Colaiácovo MP, et al. (2003) Synaptonemal Complex Assembly in C. elegans Is Dispensable for Loading Strand-Exchange Proteins but Critical for Proper Completion of Recombination. Developmental Cell 5(3):463–474.

[15] Smolikov S, et al. (2007) SYP-3 Restricts Synaptonemal Complex Assembly to Bridge Paired Chromosome Axes During Meiosis in Caenorhabditis elegans. Genetics 176(4):2015–2025.

[16] Smolikov S, Schild-Prüfert K, Colaiácovo MP (2009) A Yeast Two-Hybrid Screen for SYP-3 Interactors Identifies SYP-4, a Component Required for Synaptonemal Complex Assembly and Chiasma Formation in Caenorhabditis elegans Meiosis. PLOS Genetics 5(10):e1000669.

[17] Cahoon CK, Hawley RS (2016) Regulating the construction and demolition of the synaptonemal complex. Nature Structural & Molecular Biology 23(5):369–377.

[18] Hurlock ME, et al. (2020) Identification of novel synaptonemal complex components in C. elegans. Journal of Cell Biology 219(5):e201910043.

[19] Zhang Z, et al. (2020) Multivalent weak interactions between assembly units drive synaptonemal complex formation. Journal of Cell Biology 219(5):e201910086.

[20] Blundon JM, et al. (2024) Skp1 proteins are structural components of the synaptonemal complex in C. elegans. Science Advances 10(7):eadl4876.

[21] Fung JC, Rockmill B, Odell M, Roeder G (2004) Imposition of Crossover Interference through the Nonrandom Distribution of Synapsis Initiation Complexes. Cell 116(6):795–802.

[22] Börner G, Kleckner N, Hunter N (2004) Crossover/Noncrossover Differentiation, Synaptonemal Complex Formation, and Regulatory Surveillance at the Leptotene/Zygotene Transition of Meiosis. Cell 117(1):29–45.

[23] Capilla-Pérez L, et al. (2021) The synaptonemal complex imposes crossover interference and heterochiasmy in Arabidopsis. Proceedings of the National Academy of Sciences 118(12):e2023613118.

[24] France MG, et al. (2021) ZYP1 is required for obligate cross-over formation and crossover interference in Arabidopsis. Proceedings of the National Academy of Sciences 118(14):e2021671118.

[25] Hayashi M, Mlynarczyk-Evans S, Villeneuve AM (2010) The Synaptonemal Complex Shapes the Crossover Landscape Through Cooperative Assembly, Crossover Promotion and Crossover Inhibition During Caenorhabditis elegans Meiosis. Genetics 186(1):45–58.

[26] Libuda DE, Uzawa S, Meyer BJ, Villeneuve AM (2013) Meiotic chromosome structures constrain and respond to designation of crossover sites. Nature 502(7473):703–706.

[27] Rog O, Köhler S, Dernburg AF (2017) The synaptonemal complex has liquid crystalline properties and spatially regulates meiotic recombination factors. eLife 6:e21455.

[28] Zhang L, Köhler S, Rillo-Bohn R, Dernburg AF (2018) A compartmentalized signaling network mediates crossover control in meiosis. eLife 7:e30789.

[29] Zhang L, Stauffer W, Zwicker D, Dernburg AF (2021) Crossover patterning through kinase-regulated condensation and coarsening of recombination nodules. bioRxiv p. 2021.08.26.457865.

[30] Morgan C, et al. (2021) Diffusion-mediated HEI10 coarsening can explain meiotic crossover positioning in Arabidopsis. Nature Communications 12(1):4674.

[31] Fozard JA, Morgan C, Howard M (2023) Coarsening dynamics can explain meiotic crossover patterning in both the presence and absence of the synaptonemal complex. eLife 12:e79408.

32. [32] von Diezmann L, Bristow C, Ofer R (2024) Diffusion within the synaptonemal complex can account for signal transduction along meiotic chromosomes. bioRxiv p. 2024.05.22.595404.

[33] Durand S, et al. (2022) Joint control of meiotic crossover patterning by the synaptonemal complex and HEI10 dosage. Nature Communications 13(1):5999.

[34] MacQueen AJ (2002) Synapsis-dependent and -independent mechanisms stabilize homolog pairing during meiotic prophase in C. elegans. Genes & Development 16(18):2428–2442.

[35] de Vries FA, et al. (2005) Mouse Sycp1 functions in synaptonemal complex assembly, meiotic recombination, and XY body formation. Genes & Development 19(11):1376–1389.

[36] Espagne E, et al. (2011) Sme4 coiled-coil protein mediates synaptonemal complex assembly, recombinosome relocalization, and spindle pole body morphogenesis. Proceedings of the National Academy of Sciences 108(26):10614–10619.

[37] Köhler S, Wojcik M, Xu K, Dernbug AF (2022) Dynamic molecular architecture of the synaptonemal complex. bioRxiv p. 2020.02.16.947804.

[38] McKim KS, Peters K, Rose AM (1993) Two types of sites required for meiotic chromosome pairing in Caenorhabditis elegans. Genetics 134(3):749–768.

[39] Gordon SG, Kursel LE, Xu K, Rog O (2021) Synaptonemal Complex dimerization regulates chromosome alignment and crossover patterning in meiosis. PLOS Genetics 17(3):e1009205.

[40] Hillers KJ, Jantsch V, Martinez-Perez E, Yanowitz JL (2017) Meiosis. in WormBook. (The C. elegans Research Community).

[41] Yokoo R, et al. (2012) COSA-1 Reveals Robust Homeostasis and Separable Licensing and Reinforcement Steps Governing Meiotic Crossovers. Cell 149(1):75–87.

[42] Piñeiro López C, Rodrigues Neves AR, Čavka I, Gros OJ, Köhler S (2023) Segmentation of C. elegans germline nuclei. microPublication biology.

[43] McPeek MS, Speed TP (1995) Modeling interference in genetic recombination. Genetics 139(2):1031–1044.

[44] Broman KW, Weber JL (2000) Characterization of Human Crossover Interference. The American Journal of Human Genetics 66(6):1911–1926.

[45] Kar FM, Hochwagen A (2021) Phospho-Regulation of Meiotic Prophase. Frontiers in Cell and Developmental Biology 9:667073.

[46] Nadarajan S, et al. (2017) Polo-like kinase-dependent phosphorylation of the synaptonemal complex protein SYP-4 regulates double-strand break formation through a negative feedback loop. eLife 6:e23437.

[47] Láscarez-Lagunas LI, et al. (2022) ATM/ATR kinases link the synaptonemal complex and DNA double-strand break repair pathway choice. Current Biology 32(21):4719–4726.

[48] Kumar M, et al. (2022) The Eukaryotic Linear Motif resource: 2022 release. Nucleic Acids Research 50(D1):D497–D508.

[49] Adamo A, et al. (2008) BRC-1 acts in the inter-sister pathway of meiotic double-strand break repair. EMBO reports 9(3):287–292.

[50] Li Q, et al. (2018) The tumor suppressor BRCA1-BARD1 complex localizes to the synaptonemal complex and regulates recombination under meiotic dysfunction in Caenorhabditis elegans. PLOS Genetics 14(11):e1007701.

[51] Janisiw E, Dello Stritto MR, Jantsch V, Silva N (2018) BRCA1-BARD1 associate with the synaptonemal complex and pro-crossover factors and influence RAD-51 dynamics during Caenorhabditis elegans meiosis. PLOS Genetics 14(11):e1007653.

[52] Li Q, Hariri S, Engebrecht J (2020) Meiotic Double-Strand Break Processing and Crossover Patterning Are Regulated in a Sex-Specific Manner by BRCA1–BARD1 in Caenorhabditis elegans. Genetics 216(2):359–379.

[53] Trivedi S, Blazícková J, Silva N (2022) PARG and BRCA1–BARD1 cooperative function regulates DNA repair pathway choice during gametogenesis. Nucleic Acids Research 50(21):12291–12308.

[54] Cahoon CK, Richter CM, Dayton AE, Libuda DE (2023) Sexual dimorphic regulation of recombination by the synaptonemal complex in C. elegans. eLife 12:e84538.

[55] Woglar A, Villeneuve AM (2018) Dynamic Architecture of DNA Repair Complexes and the Synaptonemal Complex at Sites of Meiotic Recombination. Cell 173(7):1678–1691.

[56] Yang C, et al. (2022) ZYP1-mediated recruitment of PCH2 to the synaptonemal complex remodels the chromosome axis leading to crossover restriction. Nucleic Acids Research 50(22):12924–12937.

[57] Gómez-H L, et al. (2016) C14ORF39/SIX6OS1 is a constituent of the synaptonemal complex and is essential for mouse fertility. Nature Communications 7(1):13298.

[58] Kong A, et al. (2014) Common and low-frequency variants associated with genomewide recombination rate. Nature Genetics 46(1):11–16.

[59] Martin EW, Mittag T (2018) Relationship of Sequence and Phase Separation in Protein Low-Complexity Regions. Biochemistry 57(17):2478–2487.

[60] Vernon RM, et al. (2018) Pi-Pi contacts are an overlooked protein feature relevant to phase separation. eLife 7:e31486.

[61] Nott TJ, et al. (2015) Phase Transition of a Disordered Nuage Protein Generates Environmentally Responsive Membraneless Organelles. Molecular Cell 57(5):936–947.

[62] Schmidt HB, Barreau A, Rohatgi R (2019) Phase separation-deficient TDP43 remains functional in splicing. Nature Communications 10(1):4890.

[63] Swasthi HM, Basalla JL, Dudley CE, Vecchiarelli AG, Chapman MR (2023) Cell surface-localized CsgF condensate is a gatekeeper in bacterial curli subunit secretion. Nature Communications 14(1):2392.

[64] Nguyen H, Labella S, Silva N, Jantsch V, Zetka M (2018) C. elegans ZHP-4 is required at multiple distinct steps in the formation of crossovers and their transition to segregation competent chiasmata. PLOS Genetics 14(10):e1007776.

[65] Brenner S (1974) The genetics of Caenorhabditis elegans. Genetics 77(1):71–94.

[66] Paix A, Folkmann A, Seydoux G (2017) Precision genome editing using CRISPR-Cas9 and linear repair templates in C. elegans. Methods 121-122(2017):86–93.

[67] Sternberg PW, et al. (2024) WormBase 2024: status and transitioning to Alliance infrastructure. Genetics 227(1):iyae050.

[68] Frøkjær-Jensen C, et al. (2016) An Abundant Class of Non-coding DNA Can Prevent Stochastic Gene Silencing in the C. elegans Germline. Cell 166(2):343–357.

[69] Frøkjær-Jensen C, et al. (2008) Single copy insertion of transgenes in C. elegans. Nat. Genet. 40(11):1375–1383.

[70] Katoh K, Standley DM (2013) MAFFT multiple sequence alignment software version 7: Improvements in performance and usability. Molecular Biology and Evolution 30(4):772–780.

[71] Waterhouse AM, Procter JB, Martin DM, Clamp M, Barton GJ (2009) Jalview Version 2-A multiple sequence alignment editor and analysis workbench. Bioinformatics 25(9):1189–1191.

[72] McDonnell AV, Jiang T, Keating AE, Berger B (2006) Paircoil2: improved prediction of coiled coils from sequence. Bioinformatics 22(3):356–358.

[73] Phillips CM, McDonald KL, Dernburg AF (2009) Cytological Analysis of Meiosis in Caenorhabditis elegans in Meiosis: Volume 2, Cytological Methods, Methods in Molecular Biology, ed. Keeney S. (Humana Press, Totowa, NJ) Vol. 558, pp. 171–195.

[74] Köhler S, Wojcik M, Xu K, Dernburg AF (2017) Superresolution microscopy reveals the three-dimensional organization of meiotic chromosome axes in intact Caenorhabditis elegans tissue. Proceedings of the National Academy of Sciences 114(24):E4734–E4743.

[75] MacQueen AJ, et al. (2005) Chromosome Sites Play Dual Roles to Establish Homologous Synapsis during Meiosis in C. elegans. Cell 123(6):1037–1050.

[76] Schindelin J, et al. (2012) Fiji: An open-source platform for biological-image analysis. Nature Methods 9(7):676–682.

[77] Pachitariu M, Stringer C (2022) Cellpose 2.0: how to train your own model. Nature Methods 19(12):1634–1641.

[78] van der Walt S, et al. (2014) scikit-image: image processing in Python. PeerJ 2(1):e453.

[79] Horvitz HR, Chalfie M, Trent C, Sulston JE, Evans PD (1982) Serotonin and Octopamine in the Nematode Caenorhabditis elegans. Science 216(4549):1012–1014.

[80] Miller DL, Roth MB (2009) C. Elegans Are Protected from Lethal Hypoxia by an Embryonic Diapause. Current Biology 19(14):1233–1237.

[81] Ewels P, Magnusson M, Lundin S, Käller M (2016) MultiQC: summarize analysis results for multiple tools and samples in a single report. Bioinformatics 32(19):3047–3048.

[82] Bolger AM, Lohse M, Usadel B (2014) Trimmomatic: a flexible trimmer for Illumina sequence data. Bioinformatics 30(15):2114–2120.

[83] Wood DE, Lu J, Langmead B (2019) Improved metagenomic analysis with Kraken 2. Genome Biology 20(1):257.

[84] Li H (2013) Aligning sequence reads, clone sequences and assembly contigs with BWAMEM. arXiv 00(00):1–3.

[85] Danecek P, et al. (2021) Twelve years of SAMtools and BCFtools. GigaScience 10(2):1–4.

[86] Rausch T, Hsi-Yang Fritz M, Korbel JO, Benes V (2019) Alfred: interactive multi-sample BAM alignment statistics, feature counting and feature annotation for long- and short-read sequencing. Bioinformatics 35(14):2489–2491.

[87] Rausch T, Fritz MHY, Untergasser A, Benes V (2020) Tracy: basecalling, alignment, assembly and deconvolution of sanger chromatogram trace files. BMC Genomics 21(1):230.

88. [88] Porta-de-la Riva M, Fontrodona L, Villanueva A, Cerón J (2012) Basic Caenorhabditis elegans Methods: Synchronization and Observation. Journal of Visualized Experiments (64):e4019.

[89] Han M, Wei G, McManus CE, Hillier LW, Reinke V (2019) Isolated C. elegans germ nuclei exhibit distinct genomic profiles of histone modification and gene expression. BMC Genomics 20(1):500.

[90] Van Nostrand EL, et al. (2016) Robust transcriptome-wide discovery of RNA-binding protein binding sites with enhanced CLIP (eCLIP). Nature Methods 13(6):508–514.

[91] Savitski MM, et al. (2014) Tracking cancer drugs in living cells by thermal profiling of the proteome. Science 346(6205):1255784.

[92] Franken H, et al. (2015) Thermal proteome profiling for unbiased identification of direct and indirect drug targets using multiplexed quantitative mass spectrometry. Nature Protocols 10(10):1567–1593.

[93] Ahlmann-Eltze C, Huber W (2020) glmGamPoi: fitting Gamma-Poisson generalized linear models on single cell count data. Bioinformatics 36(24):5701–5702.

