## Supplemental Information for "Crossovers are regulated by a conserved and disordered synaptonemal complex domain"

Ana Rita Rodrigues Neves[1,2], Ivana Čavka[1,2], Tobias Rausch[3,4], Simone Köhler[1,*]

Ana Rita Rodrigues Neves[1,2], Ivana Čavka[1,2], Tobias Rausch[3,4], Simone Köhler[1,*]

[1] European Molecular Biology Laboratory (EMBL), Cell Biology and Biophysics Unit, Heidelberg, Germany

[2] Collaboration for joint PhD degree between EMBL and Heidelberg University, Faculty of Biosciences, Heidelberg, Germany

[3] European Molecular Biology Laboratory (EMBL), Genome Biology Unit, Heidelberg, Germany.

[4] European Molecular Biology Laboratory (EMBL), GeneCore, Heidelberg, Germany.

**A**

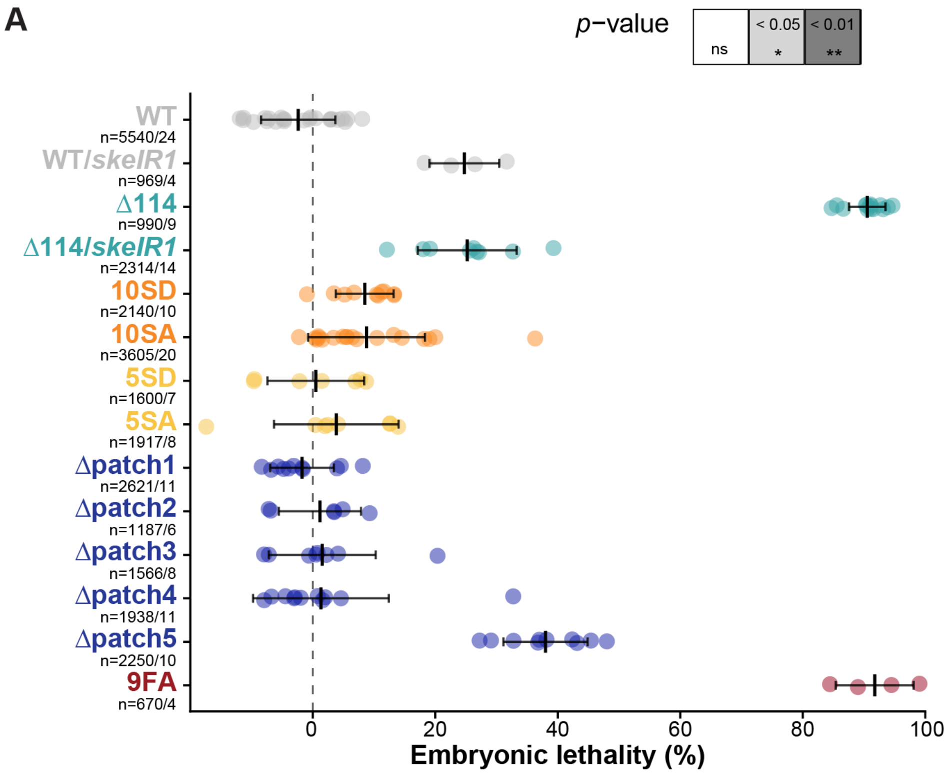


*p*−value

| | WT | WT/*skelR1* | Δ114 |
|---|---|---|---|
| WT | | $2.4 \cdot 10^{-4}$ | $3.1 \cdot 10^{-9}$ |
| WT/*skelR1* | $2.4 \cdot 10^{-4}$ | | $7.5 \cdot 10^{-4}$ |
| Δ114 | $3.1 \cdot 10^{-9}$ | $7.5 \cdot 10^{-4}$ | |
| Δ114/*skelR1* | $1.9 \cdot 10^{-7}$ | 0.94 | $6.1 \cdot 10^{-6}$ |
| 10SD | $3.6 \cdot 10^{-4}$ | | $6.4 \cdot 10^{-5}$ |
| 10SA | $5.2 \cdot 10^{-4}$ | | $1.1 \cdot 10^{-8}$ |
| 5SD | 0.40 | | $2.9 \cdot 10^{-5}$ |
| 5SA | 0.11 | | $1.2 \cdot 10^{-5}$ |
| Δpatch1 | 0.82 | | $1.7 \cdot 10^{-6}$ |
| Δpatch2 | 0.31 | | $6.4 \cdot 10^{-5}$ |
| Δpatch3 | 0.49 | | $1.2 \cdot 10^{-5}$ |
| Δpatch4 | 0.55 | | $1.7 \cdot 10^{-6}$ |
| Δpatch5 | $7.6 \cdot 10^{-8}$ | | $3.1 \cdot 10^{-6}$ |
| 9FA | $2.4 \cdot 10^{-4}$ | | 0.94 |

**B**

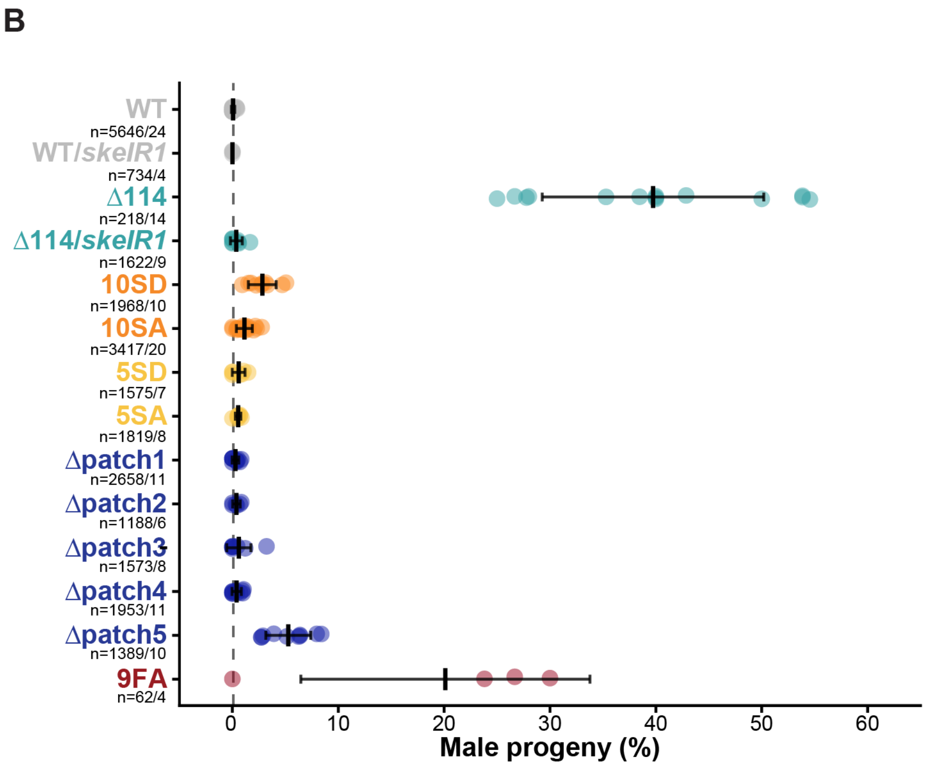


*p*−value

| | WT | WT/*skelR1* | Δ114 |
|---|---|---|---|
| WT | | 0.42 | $6.2 \cdot 10^{-7}$ |
| WT/*skelR1* | 0.42 | | $3.7 \cdot 10^{-3}$ |
| Δ114 | $6.2 \cdot 10^{-7}$ | $3.7 \cdot 10^{-3}$ | |
| Δ114/*skelR1* | $5.5 \cdot 10^{-2}$ | 0.16 | $1.6 \cdot 10^{-4}$ |
| 10SD | $1.6 \cdot 10^{-6}$ | | $1.2 \cdot 10^{-4}$ |
| 10SA | $1.1 \cdot 10^{-6}$ | | $7.8 \cdot 10^{-6}$ |
| 5SD | $1.8 \cdot 10^{-2}$ | | $4.3 \cdot 10^{-4}$ |
| 5SA | $7.1 \cdot 10^{-5}$ | | $2.5 \cdot 10^{-4}$ |
| Δpatch1 | $2.3 \cdot 10^{-2}$ | | $9.7 \cdot 10^{-5}$ |
| Δpatch2 | $1.8 \cdot 10^{-2}$ | | $7.5 \cdot 10^{-4}$ |
| Δpatch3 | 0.16 | | $2.5 \cdot 10^{-4}$ |
| Δpatch4 | $1.9 \cdot 10^{-2}$ | | $9.7 \cdot 10^{-5}$ |
| Δpatch5 | $1.6 \cdot 10^{-6}$ | | $1.2 \cdot 10^{-4}$ |
| 9FA | $8.2 \cdot 10^{-3}$ | | $2.0 \cdot 10^{-2}$ |

**Figure S1: Brood counts for strains generated in this study.**

**Figure S1: Brood counts for strains generated in this study.** (A) The quantification of embryonic lethality in the SYP-4 mutant alleles generated in this study reveals an increased lethality in *syp-4*$^{\Delta 114}$ and *syp-4*$^{9FA}$ animals with more than 90% lethality compared to 0% lethality in WT animals. Embryonic lethality is also increased in △patch5, in 10SD and 10SA animals (38%, 8% and 6%, respectively). The heterozygous balanced WT/*skeIR1* and *syp-4*$^{\Delta 114}$/*skeIR1* show a lethality of about 25% as expected. The dashed vertical line at 0% embryonic lethality corresponds to the expected value in WT animals. Error bars show mean ± standard deviations. The total number of counted eggs (E) laid by N animals is indicated for each strain as n=E/N. (B) The incidence of male progeny is significantly increased in the majority of the SYP-4 mutant alleles. This increase is more pronounced in *syp-4*$^{\Delta 114}$, *syp-4*$^{9FA}$ and △patch5 animals (40%, 20% and 5%, respectively). The dashed vertical line at 0.1% male progeny corresponds to the expected value in WT animals. Error bars show mean ± standard deviations. The total number of adult progeny (A) from N animals is indicated for each strain as n=A/N. *P*-values were calculated using the Mann-Whitney *U* test and, for comparisons to WT and *syp-4*$^{\Delta 114}$, corrected using the Benjamini-Hochberg method.

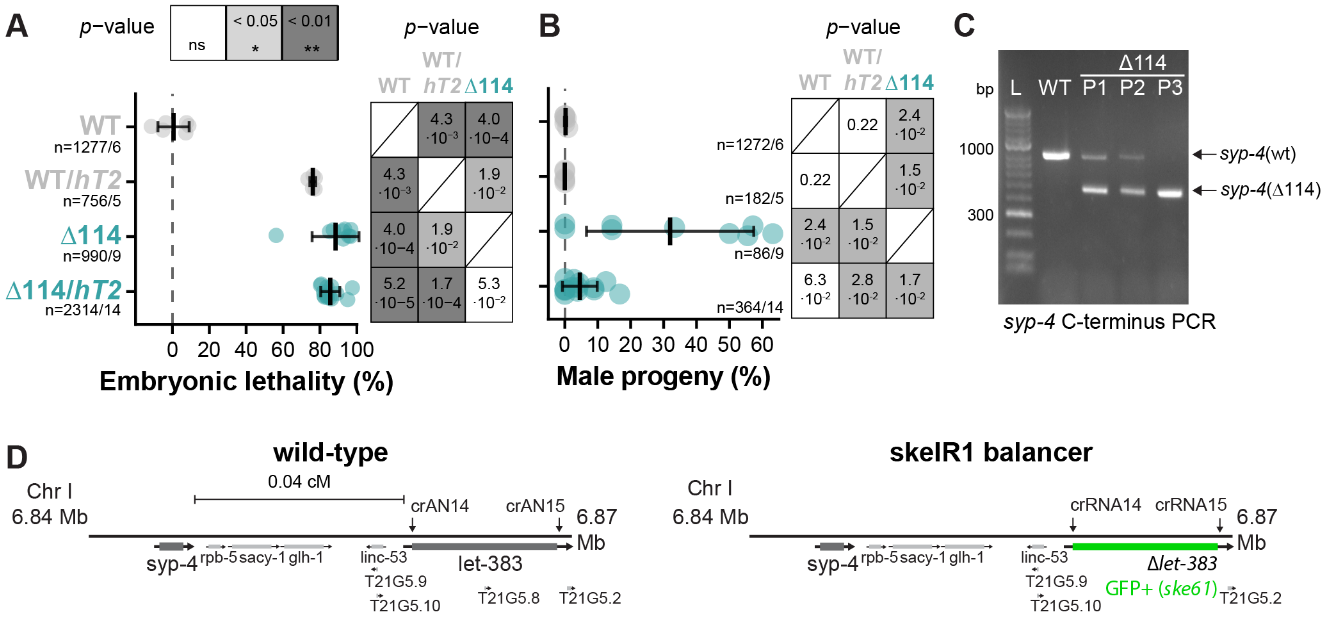


**Figure S2: The *hT2* balancer chromosome is unstable in *syp-4*$^{\Delta 114}$/*hT2* animals.** (A) The embryonic lethality in heterozygous balanced *syp-4*$^{\Delta 114}$/*hT2* animals is increased compared to the heterozygous balanced WT/*hT2* animals indicating a semi-dominant effect of the *syp-4*$^{\Delta 114}$ allele. The dashed vertical line at 0% embryonic lethality corresponds to the expected value in WT animals. Error bars show mean ± standard deviations. The total number of counted eggs (E) laid by N animals is indicated for each strain as n=E/N. (B) The incidence of male progeny is increased not only in homozygous *syp-4*$^{\Delta 114}$ animals (32%) but also heterozygous balanced *syp-4*$^{\Delta 114}$/*hT2* animals (5%) although the latter increase is not statistically significant (*p*-value=0.063). The dashed vertical line at 0.1% male progeny corresponds to the expected value in WT animals. The total number of adult progeny (A) from N animals is indicated for each strain as n=A/N. (C) Representative agarose gel for genotyping *syp-4*$^{wt}$ and *syp-4*$^{\Delta 114}$ alleles in WT animals as well as in non-green supposedly homozygous *syp-4*$^{\Delta 114}$ animals grown in the heterozygous *hT2* balancer background from three different plates (P1, P2 and P3) shows the loss of homozygosity of the *syp-4*$^{\Delta 114}$ allele with the incorporation of a *syp-4*$^{wt}$ allele in P1 and P2. *P*-values in A and B were calculated using the Mann-Whitney *U* test. Error bars show mean ± standard deviations. (D) The SYP-4-specific balancer, *ske61*, was generated in the CB4856 Hawaiian strain by replacing the *let-383* lethal gene locus with a codon-optimised GFP construct. The diagram depicts the chromosome I region between 6.84-6.78 Mbp containing the *syp-4* and *let-383* gene loci, which are 0.04 cM apart (wild-type is shown on the left). CRISPR RNAs crAN14 and crAN15 (Table S3) were used to replace the *let-383* gene with a GFP transgene.

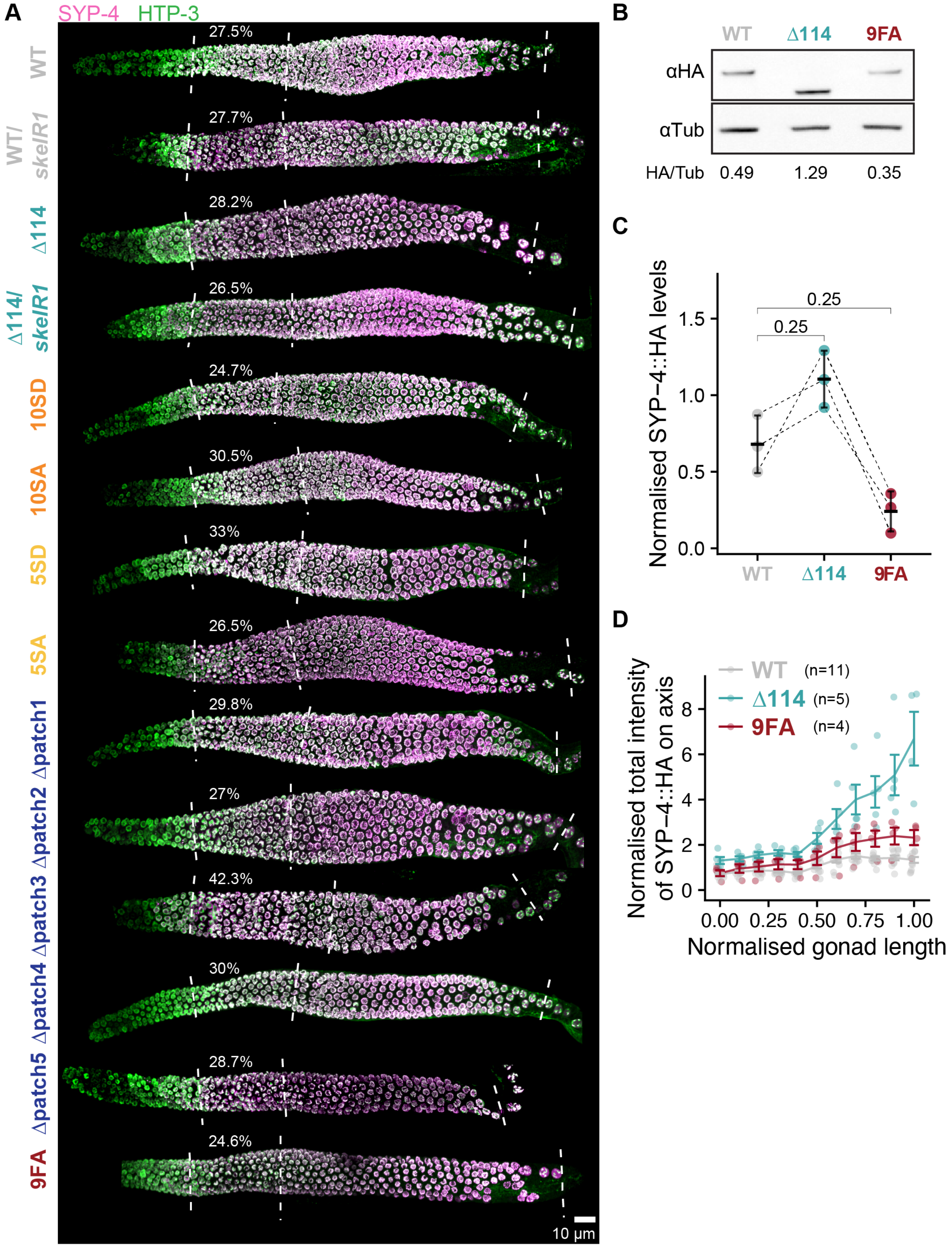


**Figure S3: SYP-4 loading is normal in *syp-4*$^{\Delta 114}$ and *syp-4*$^{9FA}$ animals.**

**Figure S3: SYP-4 loading is normal in *syp-4*$^{\Delta 114}$ and *syp-4*$^{9FA}$ animals.** (A) Representative maximum intensity projections of gonads stained for the HA-tagged synaptonemal complex protein SYP-4 (magenta) and the axis protein HTP-3 (green) used for the quantification of the length of the transition zone. The beginning and end of transition zone and pachytene are delimited by dashed lines. The percentage of transition zone compared to the combined length of transition zone and pachytene for each gonad is given. (B) Representative Western blot analysis to quantify the HA-tagged SYP-4 protein levels relative to $\alpha$-tubulin protein levels in WT, *syp-4*$^{\Delta 114}$ and *syp-4*$^{9FA}$ whole worm lysates. (C) Quantification of HA-tagged SYP-4 protein levels from Western blots in WT, *syp-4*$^{\Delta 114}$ and *syp-4*$^{9FA}$ normalised to $\alpha$-tubulin protein levels show that SYP-4 levels are slightly increased in *syp-4*$^{\Delta 114}$ animals and slightly decreased in *syp-4*$^{9FA}$ animals compared to WT animals but without statistical significance. A total of three independent experiments were performed. *P*-values were calculated using the Wilcoxon Signed-Rank test for two-paired samples. (D) Quantification of SYP-4 loading along the axis from the beginning of transition zone to the end of pachytene shows an increase in the loading of SYP-4 to the SC with meiotic progression in *syp-4*$^{\Delta 114}$ (cyan), *syp-4*$^{9FA}$ (red), and in WT (gray) animals.The amount of protein loaded in the SC is higher in *syp-4*$^{\Delta 114}$ and *syp-4*$^{9FA}$ animals than in WT animals. The normalised pachytene length was divided into 11 bins. The error bars show mean ± standard deviation.

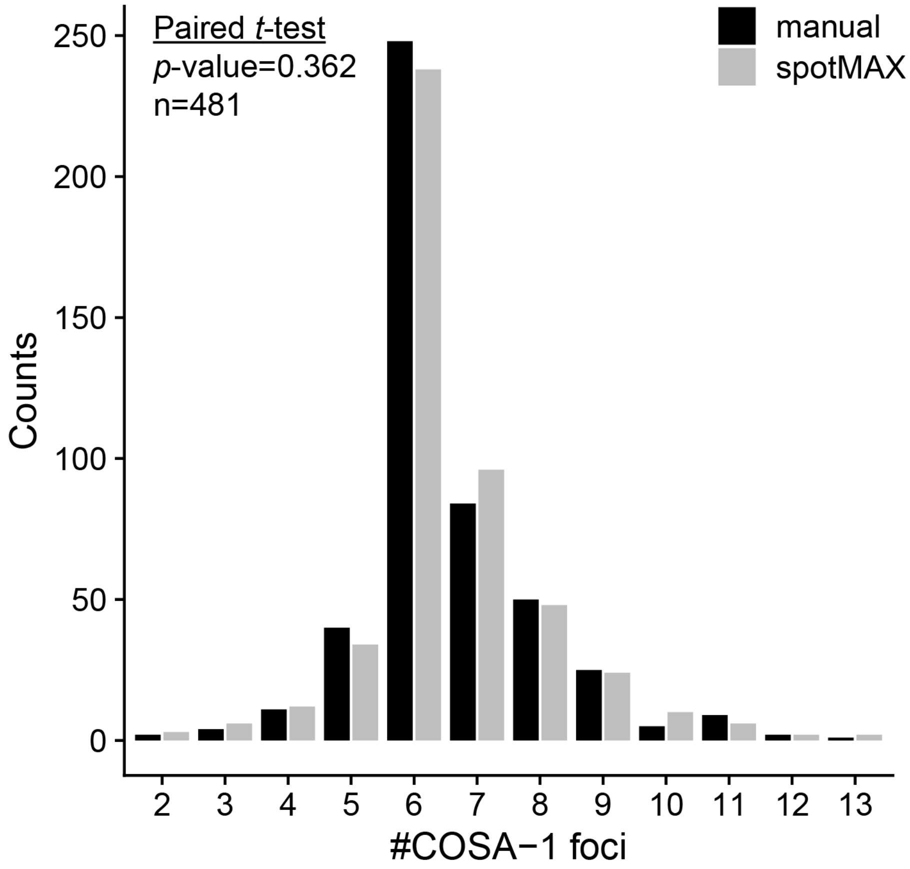


**Figure S4: spotMAX automated quantification of COSA-1 foci in 3D images from *C. elegans* fixed gonads is comparable to manual quantification.** Distribution of the number of COSA-1 quantified manually (black) and automatically using spotMAX (gray) shows no significant difference between the two methods. The same nuclei were quantified by the two approaches. A total number of 481 nuclei originating from different genetic backgrounds used in this study were quantified. No biases related to the genetic background were observed. *P*-value was calculated using the Student's *t*-test for two-paired samples.

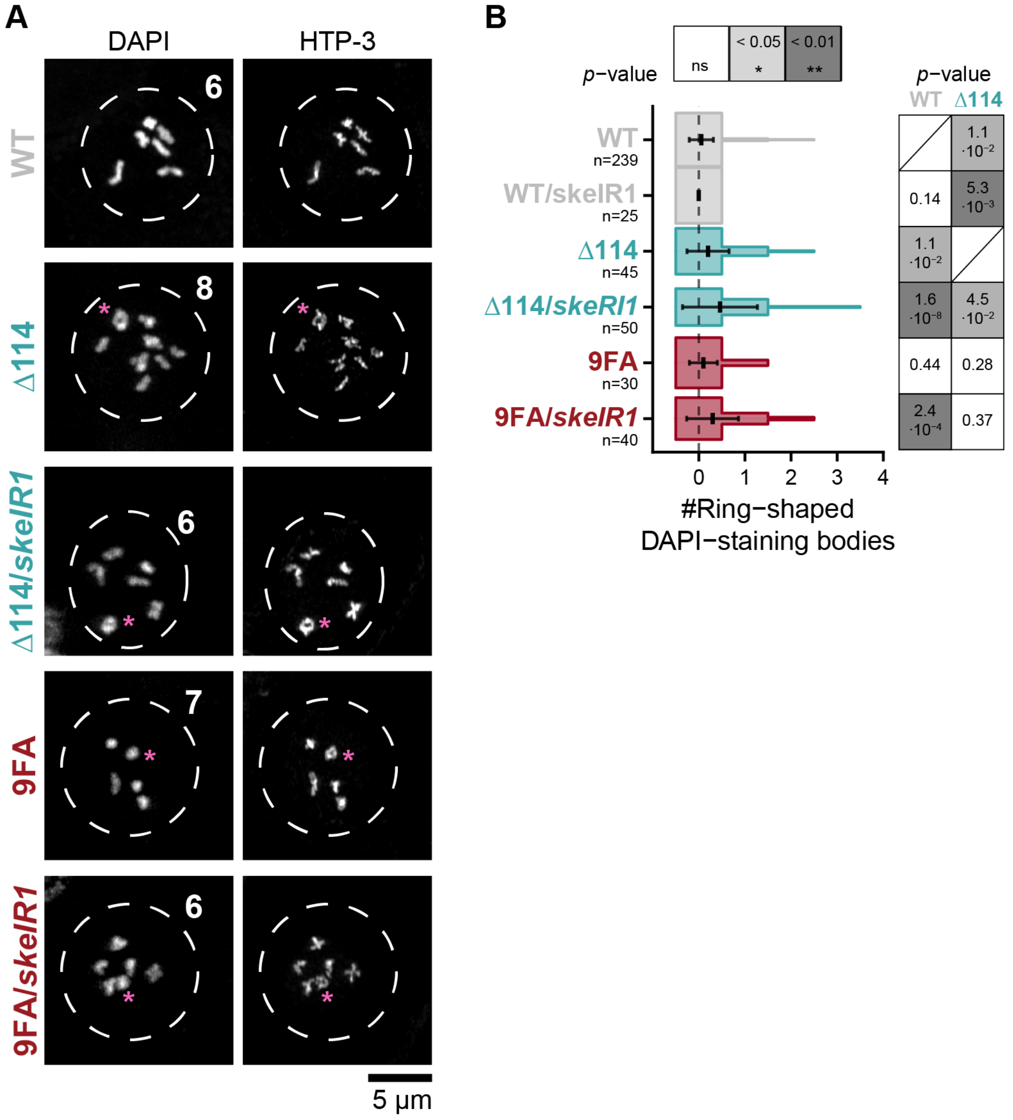


**Figure S5: Ring-shaped bivalents can be visualised during diakinesis in homozygous as well as heterozygous balanced *syp-4*$^{\Delta 114}$ and *syp-4*$^{9FA}$ animals indicating an increased number of crossovers per bivalent.** (A) Maximum intensity projections of diakinesis nuclei counterstained with DAPI (left) and stained against the axis protein HTP-3 (right). The number of DAPI-staining bodies counted in each nucleus is given. Ring-shaped DAPI-staining bodies are indicated with an asterisk (*). (B) Quantification of number of ring-shaped DAPI-staining bodies reveals ring-shaped bivalents in *syp-4*$^{\Delta 114}$ and *syp-4*$^{9FA}$ mutant alleles in both homozygosity as well as heterozygosity but not wild-type animals supporting the semi-dominant effect suggested by the number of COSA-1 foci in these mutants. The number of quantified diakinesis nuclei is given by n. *P*-values were calculated using a Gamma-Poisson generalised linear model.

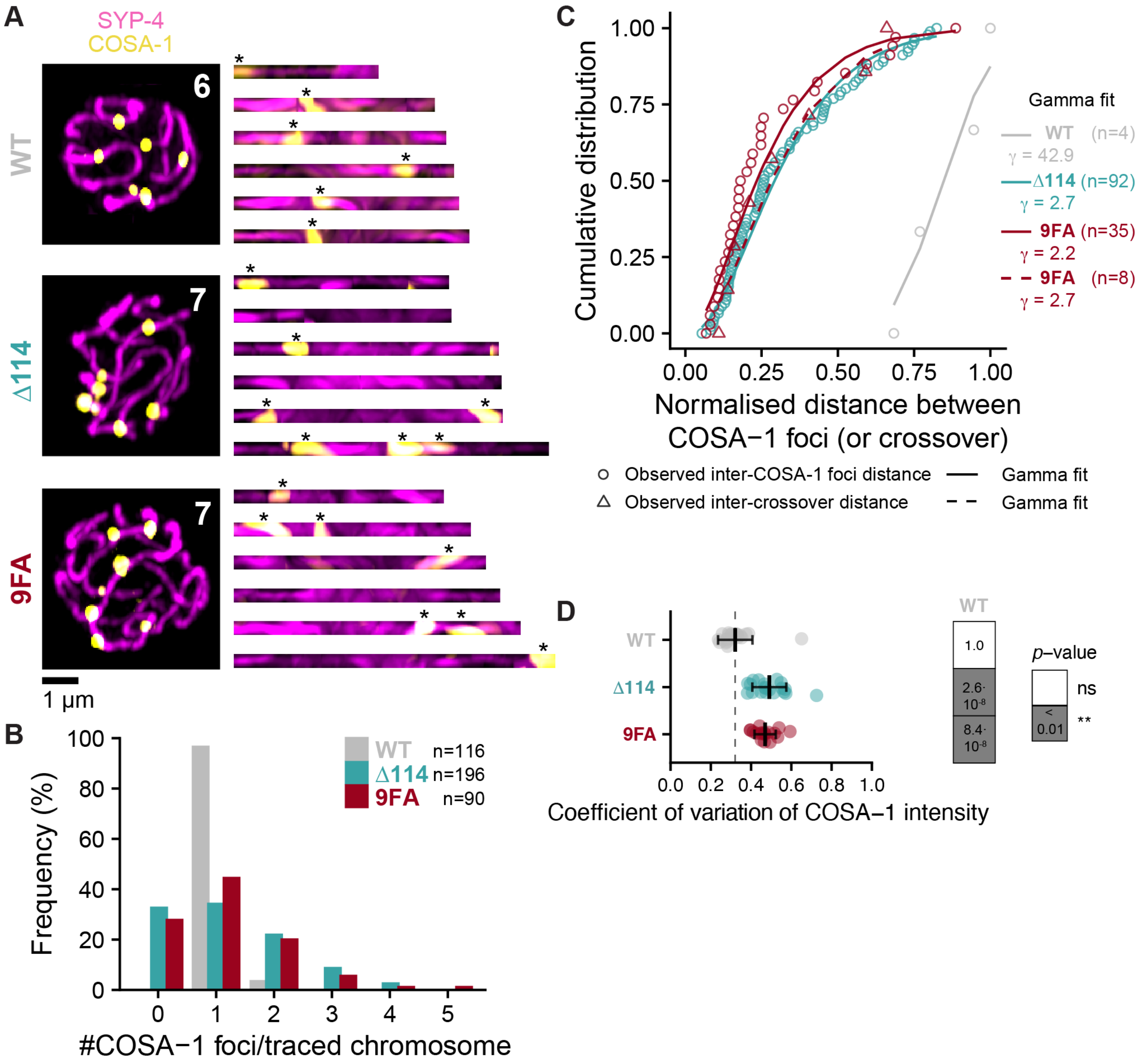

A
SYP-4
COSA-1
WT
Δ114
9FA
6
7
7
1 μm
B
Frequency (%)
#COSA−1 foci/traced chromosome
WT n=116
Δ114 n=196
9FA n=90
C
Cumulative distribution
Normalised distance between COSA−1 foci (or crossover)
Gamma fit
WT (n=4) γ = 42.9
Δ114 (n=92) γ = 2.7
9FA (n=35) γ = 2.2
9FA (n=8) γ = 2.7
Observed inter-COSA-1 foci distance
Observed inter-crossover distance
Gamma fit
Gamma fit
D
Coefficient of variation of COSA−1 intensity
WT
1.0
2.6·10-8
8.4·10-8
p–value
ns
< 0.01 **

**Figure S6: Crossover assurance and crossover interference are disrupted in *syp-4*$^{\triangle 114}$ and *syp-4*$^{9FA}$ animals.** (A) The number of COSA-1 foci per SC is tightly regulated to exactly one focus per SC in WT animals. This regulation is lost in both *syp-4*$^{9FA}$ and *syp-4*$^{\triangle 114}$ animals which have SCs with multiple or no COSA-1 foci. Maximum intensity projections of representative late pachytene nuclei stained for HA-tagged SYP-4 (magenta) and the Halo-tagged COSA-1 (yellow) are shown on the left, individual straightened SCs are shown on the right. The positions of COSA-1 foci along the chromosomes are marked by asterisks (*). (B) The distribution of the number of COSA-1 foci per traced chromosome in WT (gray), *syp-4*$^{\triangle 114}$ (cyan) and *syp-4*$^{9FA}$ (red) late pachytene nuclei shows that 32% and 28% of the chromosomes in *syp-4*$^{\triangle 114}$ and *syp-4*$^{9FA}$, respectively, have no COSA-1 focus indicating loss of assurance. At the same time, about 33% of the traced chromosomes in *syp-4*$^{\triangle 114}$ and *syp-4*$^{9FA}$ have more than one COSA-1 focus per chromosome indicating a reduction in crossover interference strength. (C) Fitting a gamma distribution to the cumulative distribution function of normalised inter-COSA-1 distances (empty circles; filled line) in WT (gray), *syp-4*$^{\triangle 114}$ (cyan) and *syp-4*$^{9FA}$ (red) indicated that crossover interference is severely reduced in *syp-4*$^{\triangle 114}$ and *syp-4*$^{9FA}$ animals with a $\gamma$ shape factor of 2.7 and 2.2, respectively, compared to a $\gamma$ factor of 42.9 in WT animals. The severe reduction of crossover interference observed in *syp-4*$^{9FA}$ was confirmed genetically by the analysis of normalised inter-crossover distances (empty triangles; dashed line) which have a $\gamma$ factor of 2.7. (D) The variability of fluorescence intensities of COSA-1 foci measured by the coefficient of variation (s.d./mean) is higher in *syp-4*$^{\triangle 114}$ and *syp-4*$^{9FA}$ than WT animals. Error bars show mean ± standard deviations. *P*-values were calculated using the Mann-Whitney *U* test.

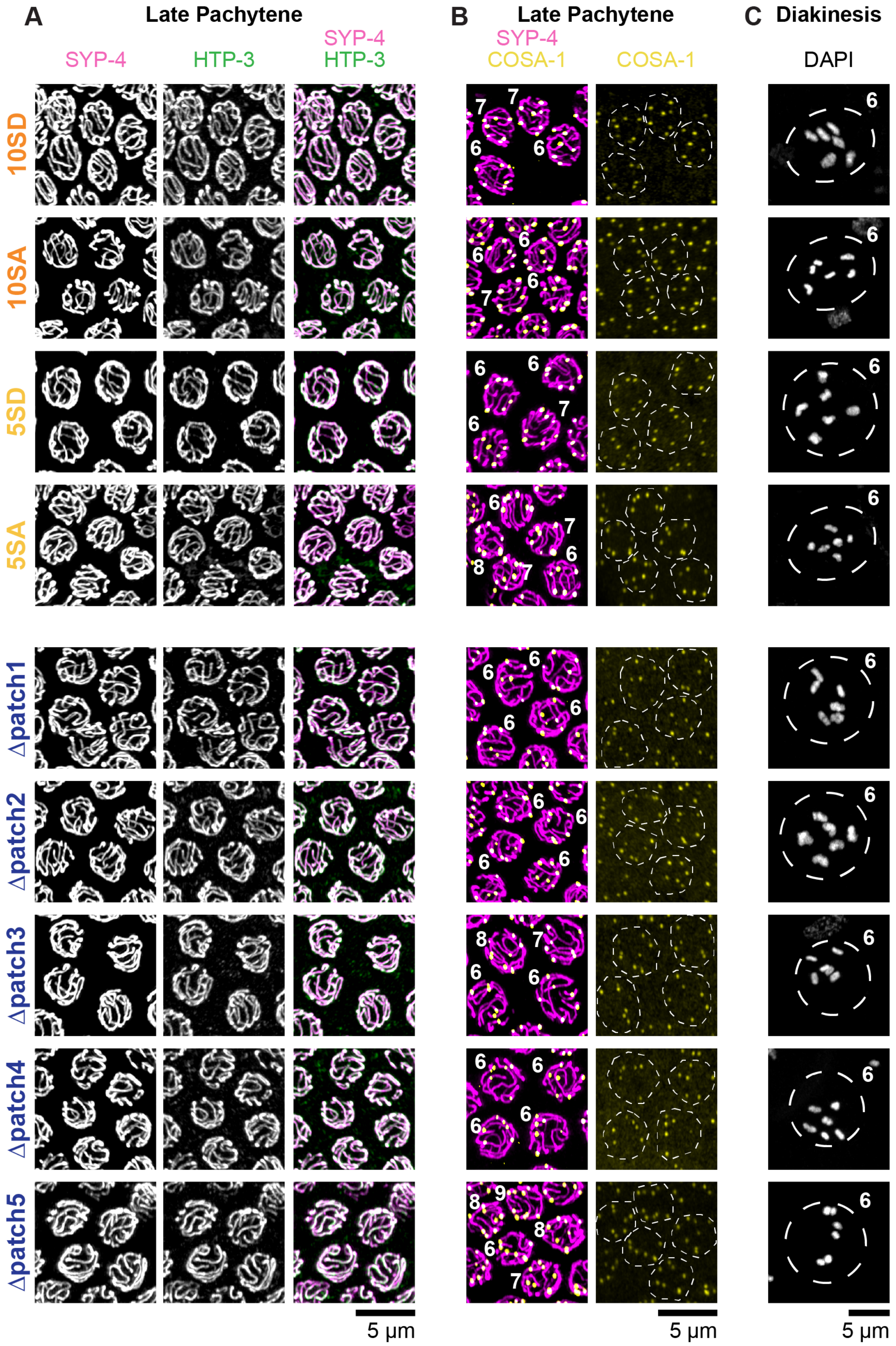


**Figure S7: Robust crossover interference but not assurance or synapsis depends on the last 32 amino acids of SYP-4 and is modulated by phosphorylation of the SYP-4 C-terminus.**

**Figure S7: Robust crossover interference but not assurance or synapsis depends on the last 32 amino acids of SYP-4 and is modulated by phosphorylation of the SYP-4 C-terminus.** (A) Maximum intensity projections of late pachytene nuclei stained for the HA-tagged synaptonemal complex protein SYP-4 (magenta, left) and the axis protein HTP-3 (green, center). The merged image is shown on the right. (B) Maximum intensity projections of late pachytene nuclei stained for the HA-tagged synaptonemal complex SYP-4 (magenta) and the Halo-tagged crossover marker COSA-1 (yellow). (C) Maximum intensity projections of diakinesis nuclei counterstained with DAPI.

**Table S1:** Total number of peptides containing each serine and threonine residue from position 440 and number of peptides containing a phosphorylated form of serine/threonine at the respective position. A total of 3 experiments were performed. * marks residues mutated in 10SD/10SA animals and †marks residues mutated in 5SD/5SA animals.

| Position | Amino Acid | #Peptides covering amino acid residue | #Peptides with phosphorylated amino acid residue |
|---|---|---|---|
| 441 | T | 14 | 0 |
| 447* | S | 12 | 4 |
| 485* | S | 10 | 5 |
| 489 | S | 10 | 0 |
| 492† | S | 15 | 0 |
| 496*† | S | 5 | 1 |
| 503 | S | 11 | 0 |
| 506 | T | 6 | 0 |
| 507 | S | 6 | 0 |
| 531* | S | 0 | 0 |
| 536* | S | 0 | 0 |
| 537* | T | 0 | 0 |
| 554* | S | 17 | 5 |
| 557 | T | 17 | 0 |
| 559† | S | 17 | 0 |
| 566* | S | 0 | 0 |
| 578*† | S | 0 | 0 |
| 592 | T | 5 | 0 |
| 593* | S | 5 | 0 |
| 599† | S | 5 | 0 |
| 600 | T | 5 | 0 |
| 601 | S | 5 | 0 |

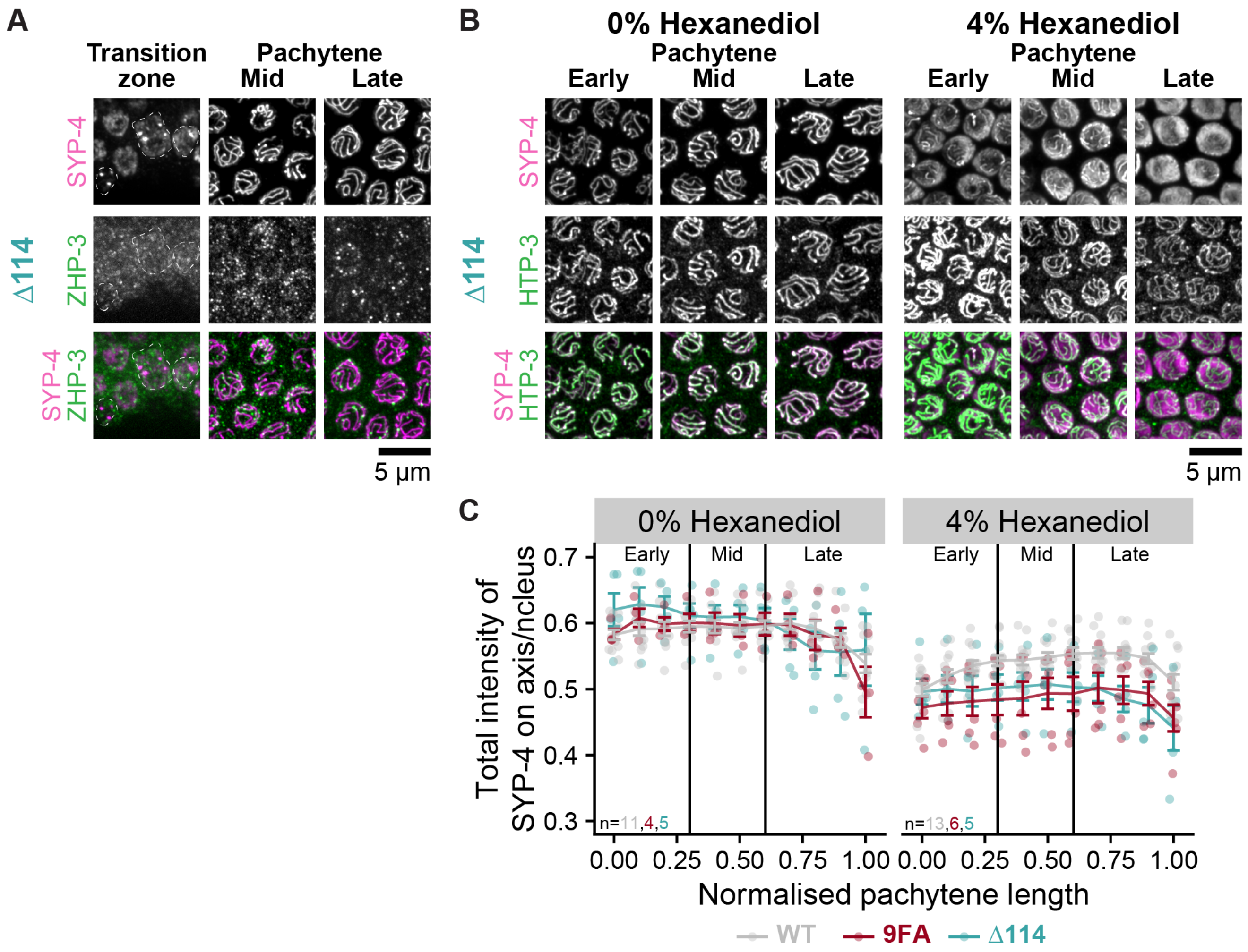


**Figure S8: The crossover formation defects observed in *syp-4*$^{\Delta 114}$ animals are accompanied by a change in the biophysical properties of the SC and the mislocalisation of ZHP-3.** (A) Maximum intensity projections of transition zone, mid and late pachytene nuclei stained for the HA-tagged synaptonemal complex protein SYP-4 (magenta, top) and the V5-tagged E3 RING ligase protein ZHP-3 (green, middle) show that ZHP-3 fails to co-localise with the SC in *syp-4*$^{\Delta 114}$ animals. The merged image is shown at the bottom. Nuclei containing putative polycomplexes during transition zone are encircled with a dashed line. (B) Maximum intensity projections of early, mid and late pachytene nuclei from extruded gonads treated with either 0% (w/v) or 4% (w/v) 1,6-hexanediol and stained for the HA-tagged synaptonemal complex protein SYP-4 (magenta, top) and the axis protein HTP-3 (green, middle). The merged image is shown at the bottom. (C) SCs in *syp-4*$^{9FA}$ and *syp-4*$^{\Delta 114}$ animals are more sensitive to 1,6-hexanediol than SCs of WT animals. The ratio of the total SYP-4 intensity on the axis to total SYP-4 intensity per nucleus in WT (gray), *syp-4*$^{9FA}$ (red) and *syp-4*$^{\Delta 114}$ (cyan) are shown during pachytene. Axis/nucleus ratios are shown along the gonad dividing pachytene into 11 bins.

**Table S2:** List of *C. elegans* strains used in this study.

| Strain Code | Genotype | Source | Referred to as |
|---|---|---|---|
| N2 | wild-type | CGC | N2 |
| CB4856 | wild-type | CGC | - |
| SMN440 | *syp-4(ie29[syp-4::ha])* I | This study | WT (in Fig. 4B and S3D) |
| SMN251 | *syp-4(ie29[syp-4::ha])* I; *cosa-1(ske25[halo::cosa-1])* III | This study | WT (all remaining Figures) |
| SMN378 | *syp-4(ie29[syp-4::ha])/skeIR1(ske61[let-383 + eft-3p::gfp::NLS::tbb-2 3'UTR]*, CB4856>N2*])* I; *cosa-1(ske25[halo::cosa-1])* III | This study | WT/*skeIR1* |
| SMN262 | *syp-4(ske35-1[syp-4(NP)::ha])* I; *cosa-1(ske25[halo::cosa-1])* III | This study | WT (in Fig. S2) |
| SMN263 | *syp-4(ie29[syp-4::ha])* I/*hT2 [bli-4(e937) let-?(q782) qIs48]* (I;III); *cosa-1(ske25[halo::cosa-1])* III | This study | WT/*hT2* |
| SMN426 | *syp-4(ie29[syp-4::ha]) zhp-3(ske71-1[zhp-3::v5]* I; *cosa-1(ske25[halo::cosa-1])* III | This study | WT (in Fig. 4A) |
| SMN242 | *syp-4(ske19-2[syp-4(D114)::ha])* I/*hT2 [bli-4(e937) let-?(q782) qIs48]* (I;III); *cosa-1(ske25[halo::cosa-1])* III | This study | Δ114 (homozygous) or Δ114/*hT2* (heterozygous) (in Fig. S3) |
| SMN380 | *syp-4(ske19-4[syp-4(D114)::ha])/skeIR1(ske61[let-383 + eft-3p::gfp::NLS::tbb-2 3'UTR*, CB4856>N2]) I; *cosa-1(ske25[halo::cosa-1])* III | This study | Δ114 (homozygous) or Δ114/*skeIR1* (heterozygous) (all remaining Figures) |
| SMN438-439 | *syp-4(ske19-4[syp-4(D114)::ha])/skeIR1(ske61[let-383 + eft-3p::gfp::NLS::tbb-2 3'UTR*, CB4856>N2]) *zhp-3(ske71-2/-3[zhp-3::v5])* I; *cosa-1(ske25[halo::cosa-1])* III | This study | Δ114 (homozygous) (in Fig. S8A) |
| SMN421 | *syp-4(ske39-1[syp-4(9FA)::ha])/skeIR1(ske61[let-383 + eft-3p::gfp::NLS::tbb-2 3'UTR*, CB4856>N2]) I; *cosa-1(ske25[halo::cosa-1])* III | This study | 9FA (homozygous) or 9FA/*skeIR1* (heterozygous) (all remaining Figures) |
| SMN428-430 | *syp-4(ske39-1[syp-4(9FA)::ha])/skeIR1(ske61[let-383 + eft-3p::gfp::NLS::tbb-2 3'UTR]*, CB4856>N2]) *zhp-3(ske71-4/-5/-6[zhp-3::v5])* I; *cosa-1(ske25[halo::cosa-1])* III | This study | 9FA (homozygous) (in Fig. 4A) |
| SMN464-465 | CB4856; *syp-4(ske39-2[syp-4(9FA)::ha])/ske61[let-383 + eft-3p::gfp::NLS::tbb-2 3'UTR]* I | This study | - |
| SMN292-293 | *syp-4(ske36-1/-2[syp-4(10D)::ha])* I; *cosa-1(ske25[halo::cosa-1])* III | This study | 10SD |
| SMN294-298 | *syp-4(ske30-1/-2/-3/-4[syp-4(10A)::ha])* I; *cosa-1(ske25[halo::cosa-1])* III | This study | 10SA |
| SMN344 | *syp-4(ske37[syp-4(5D)::ha])* I; *cosa-1(ske25[halo::cosa-1])* III | This study | 5SD |
| SMN335 | *syp-4(ske38[syp-4(5A)::ha])* I; *cosa-1(ske25[halo::cosa-1])* III | This study | 5SA |
| SMN337-339 | *syp-4(ske40-1/-2/-3[syp-4(Δ485-504aa)::ha])* I; *cosa-1(ske25[halo::cosa-1])* III | This study | Δpatch1 |
| SMN340-341 | *syp-4(ske41-1/-2[syp-4(Δ514-521aa)::ha])* I; *cosa-1(ske25[halo::cosa-1])* III | This study | Δpatch2 |
| SMN346 | *syp-4(ske42-2[syp-4(Δ527-543aa)::ha])* I; *cosa-1(ske25[halo::cosa-1])* III | This study | Δpatch3 |
| SMN342-343 | *syp-4(ske43-1/-2[syp-4(Δ559-564aa)::ha])* I; *cosa-1(ske25[halo::cosa-1])* III | This study | Δpatch4 |
| SMN461-463 | *syp-4(ske44[syp-4(Δ573-605aa)::ha])* I; *cosa-1(ske25[halo::cosa-1])* III | This study | Δpatch5 |

**Table S3:** List of CRISPR RNA sequences used to generate new alleles.

| **Allele** | **Target Gene/Allele** | **crRNA1 Name** | **crRNA1 Sequence (5'->3')** | **crRNA2 Name** | **crRNA2 Sequence (5'->3')** |
|---|---|---|---|---|---|
| *ske35[syp-4(NP)::ha]* | *ie29[syp-4::ha]* | crAN7 | ggagcaggctgcactctccc | - | - |
| *ske19[syp-4(D114)::ha]* | *ie29[syp-4::ha]* | crAN10 | agagaagttattgcttttgt | crSKie29-61 | cataatctgggacatcgtag |
| *ske30[syp-4(10A)::ha]* | *ske35[syp-4(NP)::ha]* | crAN13 | ttcctgctcttggctcaggg | crSKie29-61 | cataatctgggacatcgtag |
| *ske36[syp-4(10D)::ha]* | *ske35[syp-4(NP)::ha]* | crAN13 | ttcctgctcttggctcaggg | crSKie29-61 | cataatctgggacatcgtag |
| *ske37[syp-4(5D)::ha]* | *ske35[syp-4(NP)::ha]* | crAN13 | ttcctgctcttggctcaggg | crSKie29-61 | cataatctgggacatcgtag |
| *ske38[syp-4(5A)::ha]* | *ske35[syp-4(NP)::ha]* | crAN13 | ttcctgctcttggctcaggg | crSKie29-61 | cataatctgggacatcgtag |
| *ske39[syp-4(9FA)::ha]* | *ske35[syp-4(NP)::ha]* | crAN13 | ttcctgctcttggctcaggg | crSKie29-61 | cataatctgggacatcgtag |
| *ske40[syp-4(Δ485-504aa)::ha]* | *ske35[syp-4(NP)::ha]* | crAN13 | ttcctgctcttggctcaggg | crSKie29-61 | cataatctgggacatcgtag |
| *ske41[syp-4(Δ514-521aa)::ha]* | *ske35[syp-4(NP)::ha]* | crAN13 | ttcctgctcttggctcaggg | crSKie29-61 | cataatctgggacatcgtag |
| *ske42[syp-4(Δ527-543aa)::ha]* | *ske35[syp-4(NP)::ha]* | crAN13 | ttcctgctcttggctcaggg | crSKie29-61 | cataatctgggacatcgtag |
| *ske43[syp-4(Δ559-564aa)::ha]* | *ske35[syp-4(NP)::ha]* | crAN13 | ttcctgctcttggctcaggg | crSKie29-61 | cataatctgggacatcgtag |
| *ske44[syp-4(Δ573-605aa)::ha]* | *ske35[syp-4(NP)::ha]* | crAN13 | ttcctgctcttggctcaggg | crSKie29-61 | cataatctgggacatcgtag |
| *ske25[halo::cosa-1]* | *cosa-1* | crSK34 | aagtgtcaatgtcaagttct | - | - |
| *ske61[let-383 + eft-3p::gfp::NLS::tbb-2 3'UTR]* | *let-383* | crAN14 | gacgtctcccgcaacatttg | crAN15 | gagttagcagctctccagtt |
| *ske71[zhp-3::v5]* | *zhp-3* | crAN17 | gagattaaaacattaatcgg | - | - |

### **Table S4:** List of template sequences used to generate new alleles.

| Allele | Gene Target | Repair Template Sequence (5’->3’) |
|---|---|---|
| *ske35[syp-4(NP)::ha]* | *ie29[syp-4::ha]* | caaggacattctagccacggagcaggccgccctgagccaagagcaggaaccagagatcgttga-<br>gaagcag |
| *ske19[syp-4(D114)::ha]* | *ie29[syp-4::ha]* | gatgtagaagacgaagaagaagtgatgagcgaaaacggtagcaacaaatatccctacgatgtccca-<br>gattatgcttagaaaattatcatg |
| *ske30[syp-4(10A)::ha]* | *ske35[syp-4(NP)::ha]* | cggtgagcaagaagaaattgagcaaccatctgtattcaaggacattctagccacggagcaggct-<br>gcgctagcccaagaacaagaaccagagatcgttgagaagcaggcagacaatgat-<br>gttcagtttgttgatggtaagtgttttacttggcaataaacattgagttattcatttcgcagatcaa-<br>caagtgggagcacaactagatgtagaagacgaagaagaagtgatggccgaaaacggatc-<br>caacaaaagcaataacttcgcctttaactttttgggaatagcaagggtacatcagcgggagaag-<br>gtggcggtgaaggcagtaagttatgaaacacgtgaaattaaaaaaaactccatttttatagacttc-<br>gactttaattttgacggaattggagcaggagatgatggtgccaacaatggtggtgccgccggagatctg-<br>gatttccttaactatgacggagaggatgaaggaaagggtgccggcaacactcaatccgatccgttcg-<br>gatttgcagccaatggtaacgcagctggtggaggtggagatggagccttcaactttaactttgacggt-<br>gacggtgaaggcggagcaactgccggagcgggcggaaacagcacctcgttctttaacttttatccatat-<br>gacgtgcccgattacgcttagaaaattatcatgtattattcagctcttgtatcatttgtatcgtttaatgtca-<br>gaaattttagattttattggaccaagaaattgcc |
| *ske36[syp-4(10D)::ha]* | *ske35[syp-4(NP)::ha]* | cggtgagcaagaagaaattgagcaaccatctgtattcaaggacattctagccacggagcaggctgctc-<br>tagaccaagaacaagaaccagagatcgttgagaagcaggcagacaatgatgttcagtttgttgatg-<br>gtaagtgttttacttggcaataaacattgagttattcatttcgcagatcaacaagtgggagcacaac-<br>tagatgtagaagacgaagaagaagtgatggacgaaaacggatccaacaaaagcaataacttc-<br>gactttaactttttgggaatagcaagggtacatcagcgggagaaggtggcggtgaaggcagtaagt-<br>tatgaaacacgtgaaattaaaaaaaactccatttttatagacttcgactttaattttgacggaattg-<br>gagcaggagatgatggtgacaacaatggtggtgacgacggagatctggatttccttaactatgacg-<br>gagaggatgaaggaaagggtgacggaaacactcaatccgatccgttcggatttgcagacaatg-<br>gtaacgcagctggtggaggtggagatggagacttcaactttaactttgacggtgacggtgaaggcg-<br>gagcaactgacggagcgggcggaaacagcacctcgttctttaacttttatccatatgacgtgcccgat-<br>tacgcttagaaaattatcatgtattattcagctcttgtatcatttgtatcgtttaatgtcagaaattttagattt-<br>tattggaccaagaaattgcc |
| *ske37[syp-4(5D)::ha]* | *ske35[syp-4(NP)::ha]* | cggtgagcaagaagaaattgagcaaccatctgtattcaaggacattctagccacggagcaggctgctc-<br>taagccaagaacaagaaccagagatcgttgagaagcaggcagacaatgatgttcagtttgttgatg-<br>gtaagtgttttacttggcaataaacattgagttattcatttcgcagatcaacaagtgggagcacaac-<br>tagatgtagaagacgaagaagaagtgatgagcgaaaacggatccaacaaagacaataacttc-<br>gactttaactttttgggaatagcaagggtacatcagcgggagaaggtggcggtgaaggcagtaagt-<br>tatgaaacacgtgaaattaaaaaaaactccatttttatagacttcgactttaattttgacggaattg-<br>gagcaggagatgatggttcgaacaatggtggttctactggagatctggatttccttaactatgacg-<br>gagaggatgaaggaaagggctcaggaaacactcaagacgatccgttcggatttgcatcaaatg-<br>gtaacgcagctggtggaggtggagatggagacttcaactttaactttgacggtgacggtgaaggcg-<br>gagcaacttcaggcgctggcggaaacgacacctcgttctttaacttttatccctacgatgtcccagattat-<br>gcttagaaaattatcatgtattattcagctcttgtatcatttgtatcgtttaatgtcagaaattttagattttattg-<br>gaccaagaaattgcc |
| *ske38[syp-4(5A)::ha]* | *ske35[syp-4(NP)::ha]* | cggtgagcaagaagaaattgagcaaccatctgtattcaaggacattctagccacggagcaggctgctc-<br>taagccaagaacaagaaccagagatcgttgagaagcaggcagacaatgatgttcagtttgttgatg-<br>gtaagtgttttacttggcaataaacattgagttattcatttcgcagatcaacaagtgggagcacaacta-<br>gatgtagaagacgaagaagaagtgatgagcgaaaacggatccaacaaagccaataacttcgc-<br>ctttaactttttgggaatagcaagggtacatcagcgggagaaggtggcggtgaaggcagtaagt-<br>tatgaaacacgtgaaattaaaaaaaactccatttttatagacttcgactttaattttgacggaattg-<br>gagcaggagatgatggttcgaacaatggtggttctactggagatctggatttccttaactatgacg-<br>gagaggatgaaggaaagggctcaggaaacactcaagccgatccgttcggatttgcatcaaatg-<br>gtaacgcagctggtggaggtggagatggagccttcaactttaactttgacggtgacggtgaaggcg-<br>gagcaacttcaggcgctggcggaaacgccacctcgttctttaacttttatccctacgatgtcccagattat-<br>gcttagaaaattatcatgtattattcagctcttgtatcatttgtatcgtttaatgtcagaaattttagattttattg-<br>gaccaagaaattgcc |

*Continued on next page*

Table S4 – *Continued from previous page*

| Allele | Target Gene/Allele | Repair Template Sequence (5’->3’) |
|---|---|---|
| *ske39[syp-4(9FA)::ha]* | *ske35[syp-4(NP)::ha]* | cggtgagcaagaagaaattgagcaaccatctgtattcaaggacattctagccacggagcaggctgctc-<br>taagccaagaacaagaaccagagatcgttgagaagcaggcagacaatgatgttcagtttgttgatg-<br>gtaagtgttttacttggcaataaacattgagttattcatttcgcagatcaacaagtgggagcacaacta-<br>gatgtagaagacgaagaagaagtgatgagcgaaaacggatccaacaaaagcaataacttctctgc-<br>caacgccgccgggaatagcaagggtacatcagcgggagaaggtggcggtgaaggcagtaagt-<br>tatgaaacacgtgaaattaaaaaaaactccatttttatagacttcgacgccaatgccgacggaattg-<br>gagcaggagatgatggttcgaacaatggtggttctactggagatctggatgcccttaactatgacg-<br>gagaggatgaaggaaagggctcaggaaacactcaatccgatccgttcggatttgcatcaaatg-<br>gtaacgcagctggtggaggtggagatggatcgttcaactttaactttgacggtgacggtgaaggcggag-<br>caacttcaggcgctggcggaaacagcacctcggccgccaacgcctatccctacgatgtcccagattat-<br>gcttagaaaattatcatgtattattcagctcttgtatcatttgtatcgtttaatgtcagaaattttagattttattg-<br>gaccaagaaattgcc |
| *ske40[syp-4(Δ485-504aa)::ha]* | *ske35[syp-4(NP)::ha]* | cggtgagcaagaagaaattgagcaaccatctgtattcaaggacattctagccacggagcaggctgctc-<br>taagccaagaacaagaaccagagatcgttgagaagcaggcagacaatgatgttcagtttgttgatg-<br>gtaagtgttttacttggcaataaacattgagttattcatttcgcagatcaacaagtgggagcacaac-<br>tagatgtagaagacgaagaagaagtgatgggtacatcagcgggagaaggtggcggtgaag-<br>gcagtaagttatgaaacacgtgaaattaaaaaaaactccatttttatagacttcgactttaattttgacg-<br>gaattggagcaggagatgatggttcgaacaatggtggttctactggagatctggatttccttaactat-<br>gacggagaggatgaaggaaagggctcaggaaacactcaatccgatccgttcggatttgcatcaaatg-<br>gtaacgcagctggtggaggtggagatggatcgttcaactttaactttgacggtgacggtgaaggcggag-<br>caacttcaggcgctggcggaaacagcacctcgttctttaacttttatccctacgatgtcccagattatgctta-<br>gaaaattatcatgtattattcagctcttgtatcatttgtatcgtttaatgtcagaaattttagattttattggaccaa-<br>gaaattgcc |
| *ske41[syp-4(Δ514-521aa)::ha]* | *ske35[syp-4(NP)::ha]* | cggtgagcaagaagaaattgagcaaccatctgtattcaaggacattctagccacggagcaggctgctc-<br>taagccaagaacaagaaccagagatcgttgagaagcaggcagacaatgatgttcagtttgttgatg-<br>gtaagtgttttacttggcaataaacattgagttattcatttcgcagatcaacaagtgggagcacaacta-<br>gatgtagaagacgaagaagaagtgatgagcgaaaacggatccaacaaaagcaataacttctcttt-<br>taactttttgggaatagcaagggtacatcagcgggagaaggtggcggtgtaagttatgaaacacgt-<br>gaaattaaaaaaaactccatttttataggacggaattggagcaggagatgatggttcgaacaatggtg-<br>gttctactggagatctggatttccttaactatgacggagaggatgaaggaaagggctcaggaaacact-<br>caatccgatccgttcggatttgcatcaaatggtaacgcagctggtggaggtggagatggatcgttcaactt-<br>taactttgacggtgacggtgaaggcggagcaacttcaggcgctggcggaaacagcacctcgttctt-<br>taacttttatccctacgatgtcccagattatgcttagaaaattatcatgtattattcagctcttgtatcatttg-<br>tatcgtttaatgtcagaaattttagattttattggaccaagaaattgcc |
| *ske42[syp-4(Δ527-543aa)::ha]* | *ske35[syp-4(NP)::ha]* | cggtgagcaagaagaaattgagcaaccatctgtattcaaggacattctagccacggagcaggctgctc-<br>taagccaagaacaagaaccagagatcgttgagaagcaggcagacaatgatgttcagtttgttgatg-<br>gtaagtgttttacttggcaataaacattgagttattcatttcgcagatcaacaagtgggagcacaacta-<br>gatgtagaagacgaagaagaagtgatgagcgaaaacggatccaacaaaagcaataacttctcttt-<br>taactttttgggaatagcaagggtacatcagcgggagaaggtggcggtgaaggcagtaagttat-<br>gaaacacgtgaaattaaaaaaaactccatttttatagacttcgactttaattttgacggaattggag-<br>caaactatgacggagaggatgaaggaaagggctcaggaaacactcaatccgatccgttcggatttg-<br>catcaaatggtaacgcagctggtggaggtggagatggatcgttcaactttaactttgacggtgacggt-<br>gaaggcggagcaacttcaggcgctggcggaaacagcacctcgttctttaacttttatccctacgatgtc-<br>ccagattatgcttagaaaattatcatgtattattcagctcttgtatcatttgtatcgtttaatgtcagaaattta-<br>gattttattggaccaagaaattgcc |
| *ske43[syp-4(Δ559-564aa)::ha]* | *ske35[syp-4(NP)::ha]* | cggtgagcaagaagaaattgagcaaccatctgtattcaaggacattctagccacggagcaggctgctc-<br>taagccaagaacaagaaccagagatcgttgagaagcaggcagacaatgatgttcagtttgttgatg-<br>gtaagtgttttacttggcaataaacattgagttattcatttcgcagatcaacaagtgggagcacaactagat-<br>gtagaagacgaagaagaagtgatgagcgaaaacggatccaacaaaagcaataacttctcttt-<br>taactttttgggaatagcaagggtacatcagcgggagaaggtggcggtgaaggcagtaagt-<br>tatgaaacacgtgaaattaaaaaaaactccatttttatagacttcgactttaattttgacggaattg-<br>gagcaggagatgatggttcgaacaatggtggttctactggagatctggatttccttaactatgacg-<br>gagaggatgaaggaaagggctcaggaaacactcaagcatcaaatggtaacgcagctggtggag-<br>gtggagatggatcgttcaactttaactttgacggtgacggtgaaggcggagcaacttcaggcgctggcg-<br>gaaacagcacctcgttctttaacttttatccctacgatgtcccagattatgcttagaaaattatcatgtat-<br>tattcagctcttgtatcatttgtatcgtttaatgtcagaaattttagattttattggaccaagaaattgcc |

*Continued on next page*

Table S4 – *Continued from previous page*

| **Allele** | **Target Gene/Allele** | **Repair Template Sequence (5’->3’)** |
|---|---|---|
| *ske44[syp-4(Δ573-605aa)::ha]* | *ske35[syp-4(NP)::ha]* | cggtgagcaagaagaaattgagcaaccatctgtattcaaggacattctagccacggagcaggctgctc-<br>taagccaagaacaagaaccagagatcgttgagaagcaggcagacaatgatgttcagtttgttgatg-<br>gtaagtgttttacttggcaataaacattgagttattcatttcgcagatcaacaagtgggagcacaactagat-<br>gtagaagacgaagaagaagtgatgagcgaaaacggatccaacaaaagcaataacttctcttt-<br>taactttttgggaatagcaagggtacatcagcgggagaaggtggcggtgaaggcagtaagt-<br>tatgaaacacgtgaaattaaaaaaaactccatttttatagacttcgactttaattttgacggaattg-<br>gagcaggagatgatggttcgaacaatggtggttctactggagatctggatttccttaactatgacg-<br>gagaggatgaaggaaagggctcaggaaacactcaatccgatccgttcggatttgcatcaaatg-<br>gtaacgcagctggttatccctacgatgtcccagattatgcttagaaaattatcatgtattattcagctcttgtat-<br>catttgtatcgtttaatgtcagaaattttagattttattggaccaagaaattgcc |
| *ske25[halo::cosa-1]* | *cosa-1* | (btn)cagtgaaaaatcgtgaaaactgaactgaagtgtcaatggccgagatcggaaccggattcc-<br>cattcgacccacactacgtcgaggtccttggagagcgcatgcactacgtcgacgtcggaccacgc-<br>gacggaaccccagtccttttccttcacggaaacccaacctcctcctacgtctggcgcaacatcatccca-<br>cacgtcgccccaacccaccgctgcatcgccccagaccttatcggaatgggaaagtccgacaagcca-<br>gaccttggatacttcttcgacgaccacgtccgtttcatggacgccttcatcgaggcccttggacttgag-<br>gaggtcgtccttgtcatccacgactggggatccgcccttggattccactgggccaagcgcaaccca-<br>gagcgcgtcaagggaatcgccttcatggagttcatccgcccaatcccaacctgggacgagtggcca-<br>gagttcgcccgcgagaccttccaagccttccgcaccaccgacgtcggacgtaagcttatcatcgac-<br>caaaacgtcttcatcgagggaacccttccaatgggagtcgtccgtccacttaccgaggtcgagatggac-<br>cactaccgcgagccattccttaacccagtcgaccgcgagccactttggcgcttcccaaacgagcttc-<br>caatcgccggagagccagccaacatcgtcgcccttgtcgaggagtacatggactggcttcaccaatc-<br>cccagtcccaaagcttcttttctggggaaccccaggagtccttatcccaccagccgaggccgcc-<br>cgtcttgccaagtcccttccaaactgcaaggccgtcgacatcggaccaggacttaaccttcttcaagag-<br>gacaacccagaccttatcggatccgagatcgcccgttggctttccacccttgagatctccggaggag-<br>gaggaagttcttcacggtgagttgtcgtttcaaaataaaatgcgaacactgc |

*Continued on next page*

Table S4 – *Continued from previous page*

| **Allele** | **Target Gene/Allele** | **Repair Template Sequence (5’->3’)** |
| --- | --- | --- |
| *ske61[let-383 + eft-3p::gfp::NLS::tbb-2 3’UTR]* | *let-383* | cttttttccgccagcacggacgtctcccgcaacatgcacctttggtcttttattgtcaacttccattggttcttc-<br>cattgtttctgttaaattaatgaatttttcataaaataaagacattatacaatataaaaatgaagaatt-<br>tattgaaaataaactgccagagagaaaaagtatgcaacactcccgccgagagtgtttgaaatggtg-<br>tacggtacattttcgtgctaggagttagatgtgcaggcagcaacgagagggggagagattttttgggc-<br>cttgtgaaattaacgtgagttttctggtcatctgactaatcatgttggtttttgttggtttattttgtttttatctttgtttt-<br>tatccagattaggaaatttaaattttatgaatttataatgaggtcaaacattcagtcccagcgtttttcct-<br>gttctcactgtttagtcgaatttttattttaggctttcaacaaatgttctaactgtcttatttgtgacctcactttt-<br>tatattttttaatttttaaaaatattagaagtttctaggataatttttcgacttttattctctctaccgtccg-<br>cactcttcttacttttaaattaaattgtttttttttcagttgggaaacactttgctcactccgtagcagccatg-<br>gcaagtttgtacaaaaaagcaggctcgccaaagaagaagcgtaaggtgtccaaggtaagttttct-<br>tatgggaaagaaggaaaaaaccgagattttacttgaaaaattgaatttttcgcgggattttcac-<br>caaaaattgttgaatattcattatttcacgctgtaaaacaaaaaaaaaaaaaaatcaaaaac-<br>tacgttgaaatcgcgtttttaagcgaattttcttcagaattgccagattttaaccccaaattttgcagtttt-<br>taaataaaatttcacctttcggctcaaattgtagattttcctgaaaatttagtacaaaaaacaatttc-<br>ctcgtaaatttttcaaatagattttcagggagaggagctcttcaccggagtcgtcccaatcctcgtc-<br>gagctcgacggagacgtcaacggacacaagttctccgtctccggagagggagagggagacgc-<br>cacctacggaaagctcaccctcaagttcatctgcaccaccggaaagctcccagtcccatggccaac-<br>cctcgtcaccaccttctgctacggagtccaatgcttctcccgttacccagaccacatgaagcgtcac-<br>gacttcttcaagtccgccatgccagagggatacgtccaagagcgtaccatcttcttcgtaagttatgtc-<br>tacctgcctgcctaccgcctaaatttttttgtgaagtttcttcaaaaaatccagaaaaaaaaaacaattttcat-<br>acgatttttttcccttaaaattgtgaattttcatgctttttagccccaaaagtcattatttgagaaaaaaatttcat-<br>acaaaaaaagttttgagaaatacacaatttttttaaatgtaattttcaaattttcaattttcaactagaaaattca-<br>caaaacttgtaaattttggaccaaaacatttatacaattacttttttttgaatctaataactacaataacta-<br>caaatttaacattttcagaaggacgacggaaactacaagacccgtgccgaggtcaagttcgaggga-<br>gacaccctcgtcaaccgtatcgagctcaagggaatcgacttcaaggaggacggaaacatcctcg-<br>gacacaagctcgagtacaactacaactcccacaacgtctacatcatggccgacaagcaaaa-<br>gaacggaatcaaggtcaacttcaaggtaagttctttttttgaaaagtgcagtttgtagtctaattttcatt-<br>tattttctttttaaaaaacgcatcaatttttaaatattttgggacaaaatccgaaaactgtactaatttg-<br>tagtttgtaaaattaaaaaaaaaaacgcaaaaaaaatgttttcaaaaatgcttagaataaaaataaaatttc-<br>taaaaaattgaacaaaataaacatttaaaaattcaaaaagtttgaaaaaatgcaaagtttttttaat-<br>gcaaaaaatatttaatgtttaaacaaatttaaaaaatgttgtaagaatatgtttagagacagtttt-<br>taaatgttaaattgcaaaaaaaaaaaaagcaaaaaaaaaatttgaaaaatgtaacagaaaattt-<br>taattaaaaaagtcgtgttttttttcaaacctgcaattctttgtaattttatttataaagagaaacaaat-<br>caaattgtatattacgaaaaacgatactaaaattcgaaaatttgcggtttttttgcgtaaaaaatacg-<br>gttccgtaattttcagatccgtcacaacatcgaggacggatccgtccaactcgccgaccactac-<br>caacaaaacaccccaatcggagacggaccagtcctcctcccagacaaccactacctctccac-<br>ccaatccgccctctccaaggtaagttcatagattttgaaaaaaagtttaagaactgaaaaatg-<br>gaataaaaatatttaagagcatttttaaatgtaaaattaacaaaaaaagcgcctaagaatgtttc-<br>caaaacagtaaaaaaaaaggtttaaaaaaatgcaaaaaaaaaatttaagatgcttttaattaca-<br>caaaaaatgaaagtaaaaaaaaattaaaatattgaaaaatgtatatttgtttgaaaaggtacatttttcaat-<br>gcaaaaaatgtttctaaaaattaaaaaaatatgtttaaattgtgtaaagtgtaaattaaaaaaataaaat-<br>tatgaaaaagtattgtttcatactttttctaaattttggttgaaaaatttaaaattccagctttttattcacattat-<br>gtgttctaaaaatctttccccaattttcgggaccgttttccagtatatttccatttttcaaaaagatttttaact-<br>gaaattcatgttttcaatgctaaaaatcaataaaaaaagaatttttcaggacccaaacgagaagcgt-<br>gaccacatggtcctcctcgagttcgtcaccgccgccggaatcacccacggaatggacgagctcta-<br>caagtcccgtcgtcgtaaggccaacccaaccaagctctccgagaacgccaagaagctcgccaag-<br>gaggtcgagaacaacccagctttcttgtacaaagtgggataaatgcaaaatcctttcaagcattc-<br>ccttcttctctatcactcttctttctttttgtcaaaaaattctctcgctaatttatttgctttttttaatgttattattt-<br>tatgactttttatagtcactgaaaagtttgcatctgagtgaagtgaatgctatcaaaatgtgattctgtct-<br>gatgtactttcacaatctctcttcaattccattttgaagtgctttaaacccgaaaggttgagaaaaatgc-<br>gagcgctcaaatatttgtattgtgttcgttgagtgacccaacaaaaagaggaaactttattgtgccgccaa-<br>gaaaaaagtctcagttaggttatttttatatgtatatatatatatattta |
| *ske71[zhp-3::v5]* | *zhp-3* | gaaaccgatcaatggtcggagcttcattggacccgccgatggaggaaagccaattccaaacc-<br>cacttcttggactcgactccacctaatgttttaatctcgtttttttctgaattcgttc |

**Table S5:** List of primer sequences and restriction enzymes used for genotyping the new alleles. For the detection of silent mutations restriction sites were added (+) or removed (-) in the new allele.

| Allele | Forward Primer Name | Forward Primer Sequence (5'->3') | Reverse Primer Name | Reverse Primer Sequence (5'->3') | Restriction sites |
|---|---|---|---|---|---|
| *ske19[syp-4(D114)::ha]* | AN45 | gaatcaatcctagatgagcccgaag | OR236 | gaggcaatttcttggtccaataa | - |
| *ske30[syp-4(10A)::ha]* | AN45 | gaatcaatcctagatgagcccgaag | OR236 | gaggcaatttcttggtccaataa | HaeIII (-); BamHI (+); XmnI (-) |
| *ske36[syp-4(10D)::ha]* | AN45 | gaatcaatcctagatgagcccgaag | OR236 | gaggcaatttcttggtccaataa | HaeIII (-); BamHI (+); XmnI (-) |
| *ske37[syp-4(5D)::ha]* | AN45 | gaatcaatcctagatgagcccgaag | OR236 | gaggcaatttcttggtccaataa | HaeIII (-); BamHI (+); XmnI (-) |
| *ske38[syp-4(5A)::ha]* | AN45 | gaatcaatcctagatgagcccgaag | OR236 | gaggcaatttcttggtccaataa | HaeIII (-); BamHI (+); XmnI (-) |
| *ske39[syp-4(9FA)::ha]* | AN45 | gaatcaatcctagatgagcccgaag | OR236 | gaggcaatttcttggtccaataa | HaeIII (-); BamHI (+); XmnI (-) |
| *ske40[syp-4(Δ485-504aa)::ha]* | AN45 | gaatcaatcctagatgagcccgaag | OR236 | gaggcaatttcttggtccaataa | HaeIII (-); XmnI (-) |
| *ske41[syp-4(Δ514-521aa)::ha]* | AN45 | gaatcaatcctagatgagcccgaag | OR236 | gaggcaatttcttggtccaataa | HaeIII (-); BamHI (+); XmnI (-) |
| *ske42[syp-4(Δ527-543aa)::ha]* | AN45 | gaatcaatcctagatgagcccgaag | OR236 | gaggcaatttcttggtccaataa | HaeIII (-); BamHI (+); XmnI (-) |
| *ske43[syp-4(Δ559-564aa)::ha]* | AN45 | gaatcaatcctagatgagcccgaag | OR236 | gaggcaatttcttggtccaataa | HaeIII (-); BamHI (+); XmnI (-) |
| *ske44[syp-4(Δ573-605aa)::ha]* | AN45 | gaatcaatcctagatgagcccgaag | OR236 | gaggcaatttcttggtccaataa | HaeIII (-); BamHI (+); XmnI (-) |
| *ske25[halo::cosa-1]* | SK449 | gtattggtctcgcaccgaac | SK450 | ctgctgatacggcaggtgta | - |
| *ske61[let-383 + eft-3p::gfp::NLS::tbb-2 3'UTR]* | AN80_external | ctctgtccaactgtacccattc | AN82_internal | ctgcacatctaactcctagcac | - |
| *ske61[let-383 + eft-3p::gfp::NLS::tbb-2 3'UTR]* | AN85_internal | gagaagcgtgaccacatgg | AN86_external | cagatggaaatgagagaaacaggc | - |
| *ske71[zhp-3::v5]* | IC101 | ctcactctctcattccggg | IC102 | cagatgtgaactaggtagagaaaaag | - |

**Table S6:** Embryonic viability and male progeny for each of the strains used in this study. The average embryonic lethality and male progeny, and the corresponding standard deviation (SD), are given for each strain. *P*-values were calculated using the Mann-Whitney test and adjusted using the Benjamini-Hochberg method for comparing the embryonic lethality and male progeny of each strain to N2. Strains containing epitope-tagged wild-type genes are marked by a †. Statistical significance: * *p*-value<0.05, ** *p*-value<0.01, *** *p*-value<0.001.

| Strain Code | Genotype | #Parents | #Eggs | #Adults | #Males | Embryonic Lethality | | Male Progeny | |
|---|---|---|---|---|---|---|---|---|---|
| | | | | | | Average +/- SD (%) | Adjusted p-value | Average +/- SD (%) | Adjusted p-value |
| N2 | wild-type | 43 | 10229 | 10837 | 8 | -6.72 +/- 8.16 | - | 0.07 +/- 0.15 | - |
| CB4856 | wild-type | 17 | 3050 | 3121 | 2 | -1.23 +/- 21.98 | 2.58E-01 | 0.05 +/- 0.22 | 3.23E-01 |
| SMN440 † | *syp-4(ie29[syp-4::ha])* I | 7 | 1478 | 1610 | 1 | -9.2 +/- 3.89 | 2.17E-01 | 0.05 +/- 0.14 | 8.71E-01 |
| SMN251 † | *syp-4(ie29[syp-4::ha])* I; *cosa-1(ske25[halo::cosa-1])* III | 24 | 5540 | 5646 | 4 | -2.38 +/- 6.07 | 5.78E-02 | 0.07 +/- 0.16 | 9.45E-01 |
| SMN378 | *syp-4(ie29[syp-4::ha])/skeIR1(ske61[let-383 + eft-3p::gfp::NLS::tbb-2 3'UTR*, CB4856>N2]) I; *cosa-1(ske25[halo::cosa-1])* III | 4 | 969 | 734 | 0 | 24.75 +/- 5.71 | 2.06E-03 ** | 0 +/- 0 | 4.00E-01 |
| SMN262 † | *syp-4(ske35-1[syp-4(NP)::ha])* I; *cosa-1(ske25[halo::cosa-1])* III | 6 | 1277 | 1272 | 3 | 0.6 +/- 8.47 | 8.50E-02 | 0.22 +/- 0.37 | 3.23E-01 |
| SMN263 | *syp-4(ie29[syp-4::ha])* I/*hT2 [bli-4(e937) let-?(q782) qIs48]* (I;III); *cosa-1(ske25[halo::cosa-1])* III | 5 | 756 | 182 | 0 | 76.12 +/- 1.62 | 7.31E-04 *** | 0 +/- 0 | 3.55E-01 |
| SMN426 † | *syp-4(ie29[syp-4::ha]) zhp-3(ske71-1[zhp-3::v5]* I; *cosa-1(ske25[halo::cosa-1])* III | 5 | 1061 | 1206 | 7 | -13.77 +/- 10.43 | 1.71E-01 | 0.5 +/- 0.47 | 1.54E-02 * |
| SMN242 | *syp-4(ske19-2[syp-4(D114)::ha])* I; *cosa-1(ske25[halo::cosa-1])* III | 9 | 990 | 86 | 32 | 88.45 +/- 12.71 | 1.04E-05 *** | 31.97 +/- 25.39 | 5.27E-05 *** |
| SMN242 | *syp-4(ske19-2[syp-4(D114)::ha])* I/*hT2* (I;III); *cosa-1(ske25[halo::cosa-1])* III | 14 | 2314 | 364 | 18 | 85.61 +/- 5.14 | 2.03E-07 *** | 4.57 +/- 5.27 | 1.77E-04 *** |
| SMN380 | *syp-4(ske19-4[syp-4(D114)::ha])* I; *cosa-1(ske25[halo::cosa-1])* III | 14 | 2402 | 218 | 85 | 90.52 +/- 2.98 | 2.03E-07 *** | 39.74 +/- 10.46 | 2.50E-09 *** |
| SMN380 | *syp-4(ske19-4[syp-4(D114)::ha])/skeIR1(ske61[let-383 + eft-3p::gfp::NLS::tbb-2 3'UTR*, CB4856>N2]) I; *cosa-1(ske25[halo::cosa-1])* III | 9 | 2180 | 1622 | 6 | 25.23 +/- 8.08 | 1.04E-05 *** | 0.37 +/- 0.56 | 4.88E-02 * |
| SMN438-439 | *syp-4(ske19-4[syp-4(D114)::ha]) zhp-3(ske71-2/-3[zhp-3::v5])* I; *cosa-1(ske25[halo::cosa-1])* III | 4 | 394 | 53 | 24 | 87.82 +/- 2.89 | 2.06E-03 ** | 42.28 +/- 16.57 | 5.60E-05 *** |
| SMN421 | *syp-4(ske39-1[syp-4(9FA)::ha])* I; *cosa-1(ske25[halo::cosa-1])* III | 4 | 670 | 62 | 16 | 91.75 +/- 6.36 | 2.06E-03 ** | 20.12 +/- 13.65 | 4.48E-03 ** |
| SMN428-430 | *syp-4(ske39-1[syp-4(9FA)::ha]) zhp-3(ske71-4/-5/-6[zhp-3::v5])* I; *cosa-1(ske25[halo::cosa-1])* III | 16 | 3422 | 502 | 188 | 85.57 +/- 3.26 | 1.13E-07 *** | 37.39 +/- 9.09 | 1.10E-09 *** |
| SMN464-465 | CB4856; *syp-4(ske39-2[syp-4(9FA)::ha]* I | 7 | 1536 | 218 | 46 | 85.78 +/- 2.7 | 7.31E-05 *** | 21.44 +/- 4.82 | 9.19E-07 *** |
| SMN292-293 | *syp-4(ske36-1/-2[syp-4(10D)::ha])* I; *cosa-1(ske25[halo::cosa-1])* III | 10 | 2140 | 1968 | 51 | 8.49 +/- 4.71 | 2.14E-05 *** | 2.83 +/- 1.32 | 3.62E-08 *** |
| SMN294-298 | *syp-4(ske30-1/-2/-3/-4[syp-4(10A)::ha])* I; *cosa-1(ske25[halo::cosa-1])* III | 20 | 3605 | 3417 | 42 | 5.86 +/- 16.03 | 7.38E-06 *** | 1.13 +/- 0.76 | 5.28E-09 *** |
| SMN344 | *syp-4(ske37[syp-4(5D)::ha])* I; *cosa-1(ske25[halo::cosa-1])* III | 7 | 1600 | 1575 | 10 | 0.51 +/- 7.89 | 8.30E-02 | 0.59 +/- 0.6 | 9.73E-03 ** |
| SMN335 | *syp-4(ske38[syp-4(5A)::ha])* I; *cosa-1(ske25[halo::cosa-1])* III | 8 | 1917 | 1819 | 10 | 3.84 +/- 10.17 | 6.12E-03 ** | 0.54 +/- 0.25 | 1.24E-05 *** |

*Continued on next page*

Table S6 – *Continued from previous page*

| **Strain Code** | **Genotype** | **#Parents** | **#Eggs** | **#Adults** | **#Males** | **Embryonic Lethality** | | **Male Progeny** | |
|---|---|---|---|---|---|---|---|---|---|
| | | | | | | Average +/- SD (%) | Adjusted p-value | Average +/- SD (%) | Adjusted p-value |
| SMN337-339 | *syp-4(ske40-1/-2/-3[syp-4(Δ485-504aa)::ha])* I; *cosa-1(ske25[halo::cosa-1])* III | 11 | 2621 | 2658 | 8 | -1.75 +/- 5.2 | 6.40E-02 | 0.29 +/- 0.31 | 8.35E-03 ** |
| SMN340-341 | *syp-4(ske41-1/-2[syp-4(Δ514-521aa)::ha])* I; *cosa-1(ske25[halo::cosa-1])* III | 6 | 1187 | 1188 | 5 | 1.17 +/- 6.71 | 4.12E-02 * | 0.38 +/- 0.35 | 8.22E-03 ** |
| SMN346 | *syp-4(ske42-2[syp-4(Δ527-543aa)::ha])* I; *cosa-1(ske25[halo::cosa-1])* III | 8 | 1566 | 1573 | 9 | 1.54 +/- 8.73 | 2.86E-02 * | 0.61 +/- 1.14 | 1.62E-01 |
| SMN342-343 | *syp-4(ske43-1/-2[syp-4(Δ559-564aa)::ha])* I; *cosa-1(ske25[halo::cosa-1])* III | 11 | 1938 | 1953 | 8 | 1.33 +/- 11.09 | 2.86E-02 * | 0.4 +/- 0.44 | 8.35E-03 ** |
| SMN461-463 | *syp-4(ske44[syp-4(Δ573-605aa)::ha])* I; *cosa-1(ske25[halo::cosa-1])* III | 10 | 2250 | 1389 | 71 | 38 +/- 6.88 | 6.48E-06 *** | 5.28 +/- 2.13 | 3.62E-08 *** |

**Table S7:** List of *Caenorhabditis* species used for SYP-4 multiple sequence alignment.

| Species | Protein Assembly | Ortholog |
|---|---|---|
| *C. elegans* | WS285 | H27M09.3 |
| *C. becei* | WS285 | CSP29.g2226.t1 |
| *C. brenneri* | WS285 | CBN08735 |
| *C. briggsae* | WS285 | CBG04214 |
| *C. inopinata* | WS285 | Sp34_10146410.t1 |
| *C. inopinata* | WS285 | Sp34_10147300.t1 |
| *C. latens* | WS285 | FL83_17238 |
| *C. nigoni* | WS285 | Cni-syp-4 |
| *C. panamensis* | WS285 | CSP28.g9372.t1 |
| *C. remanei* | WS285 | CRE01249 |
| *C. sinica* | WS285 | Csp5_scaffold_00925.g16811.t1 |
| *C. sulstoni* | WS285 | CSP32.g1474.t1 |
| *C. sulstoni* | WS285 | CSP32.g1476.t1 |
| *C. tribulationis* | WS285 | CSP40.g15409.t1 |
| *C. tropicalis* | WS285 | Csp11.Scaffold627.g6892.t1 |
| *C. uteleia* | WS285 | CSP31.g22718.t1 |
| *C. waitukubuli* | WS285 | CSP39.g827.t1 |
| *C. zanzibari* | WS285 | CSP26.g17156.t1 |
